# Consequence and further analysis of the minor role of daylength in the floral initiation of sorghum crops

**DOI:** 10.64898/2026.06.12.731875

**Authors:** B. Clerget, M. Sidibe, K. vom Brocke, V. Raharinivo, D. Ortiz, G. Trouche

## Abstract

**Background and Aims:** Flowering time is assumed to be primarily controlled by daylength, but field observations disagree. For instance, mid-early sorghum varieties sown at the same time at tropical and temperate latitudes have similar flowering dates. We previously proposed an alternative framework integrating daily changes in sunrise and sunset times (dSR and dSS).

**Methods:** Variety trials of mid-late varieties from the Northern Hemisphere were carried out in Madagascar and in Argentina. Series of monthly sowings of 28 varieties were carried out in Mali between 2000 and 2009. Heading date and, in Mali, panicle initiation (PI) were recorded.

**Key Results:** In Madagascar and in Argentina mid-late varieties headed late or very late when sown in November and December, consistent with the higher dSR/dSS values of the southern hemisphere summer.

In Mali, the high interannual variability in PI time in sowings between November and February revealed that up to four photoperiodic responses — two quantitative and two qualitative — may coexist within a single late photoperiod sensitive variety. Additionally, the qualitative responses may be triggered by an internal dSR/dSS phasic coincidence, which is set at the onset of the photoperiod-sensitive phase.

These hypotheses were used to design a research model that incorporated only dSR and dSS cues. The model fitted the dataset well when two sets of parameters were calibrated for the first quantitative response, which covered two periods of over and under twelve hours per year. After daylength was added to the first qualitative response, the model also accurately predicted the duration to PI observed in three varieties sown at tropical and temperate latitudes.

**Conclusions:** The flowering time of sorghum crops mainly relies on dSR and dSS cues, while daylength plays a minor role. Two new hypotheses now require confirmation through further evidence, which is likely to come from genetic analyses.

## INTRODUCTION

Since sexual reproduction first emerged around 1.2 billion years ago, its timing has had to be synchronised between mates and with the climate. Solar photoperiod is the only stable cue across years and is responsible for the daily and seasonal timing adaptations of living organisms (Thomas and Vince-Prue, 1997). In animals, reproduction and migration are annual events that are strongly related to daylength and the implicit direction of its change, which distinguishes spring and autumn (Rowan 1926). Tournois (1912) showed the first experimental evidence in plants of the effect of daylength on flowering date. In trees and other perennial plants, floral induction that occurs many months (or even years, in the case of palm trees) before flowering, has long been associated with climatic cues or internal states (Wilkie *et al*. 2008). However, the discovery of genes homologous to *CONSTANS (CO)* and *FLOWERING LOCUS T (FT),* which regulate flowering in response to the photoperiod in Arabidopsis thaliana and many tree species, could modify our understanding of the interactions between development and growth during this prolonged process (Sun *et al*. 2022; Cai *et al*. 2025).

When animal and annual plant species are transported to a different latitude with different daylength dynamics, their reproductive calendar should change. However, rare observations of the same plant variety grown under optimum conditions at various latitudes, such as Chrysanthemum (Kamemoto and Nakasone 1953) and Sorghum (Curtis 1968; Clerget *et al*. 2021) have revealed unchanged calendars. In equatorial regions with stable daylengths throughout the year, mango and forest trees have two flowering or bud break peaks each year (Borchert *et al*. 2005; Ramírez *et al*. 2010). At these latitudes the flowering peaks are synchronised with the two annual peaks in the changes of sunrise and sunset time (dSR and dSS). The flowering peaks are also synchronised with the relative variation in Earth’s insolation (above the atmosphere), which exhibits dynamics similar to those of dSR/dSS (Borchert *et al*. 2015). Building on the dSR/dSS hypothesis, the daily sunrise and sunset times under the equator were reproduced in laboratory poultry rooms in Germany, where imported stonechat birds from Kenya were raised (Goymann *et al*. 2012). The annual morphological and sexual cycles of the two groups of birds in Germany and Kenya synchronised in response to the similar sunrise and sunset times when the length of the day was consistently 12 hours. In addition to synchronising flowering with their peers, annual plants adapt their life cycles to yearly weather variability. Thus, delayed germination results in a shorter vegetative phase, which reduces variation in flowering time (Curtis 1968; Miller *et al*. 1968). This mechanism has been used for long by farmers to modulate the sowing date of their crops. Studies in growth chambers showed that constant artificial daylengths linearly modulate the duration of the vegetative phase in quantitatively photoperiodic species (Major 1980). Additionally, reciprocal transfer experiments between two constant daylengths have demonstrated that, following an initial photoperiod-insensitive phase (PIP), the duration of the photoperiod-sensitive phase (PSP) until panicle initiation is linearly proportional to the duration experienced under each day length (Collinson *et al*. 1992; Ellis *et al*. 1992; Alagarswamy *et al*. 1998). These results were then used to create quantitative models that assumed a daily accumulation of progress proportional to daylength and temperature towards flowering time, and were calibrated using seasonal data from field crops (Summerfield *et al*. 1997). In short-day qualitative photoperiodic species, flowering cannot be initiated anymore when daylength exceeds a specific threshold, being delayed until the return of the same threshold when daylength decreases, as in kenaf and chickpea (Carberry *et al*. 1992, 2001).

However, monthly sowings in the field often show a more complex annual response. Tropical cereals and pulses can be sown and grown all over the year because they tolerate daily midday temperature over 40°C and relative humidity below 10% during the dry months provided they are permanently well irrigated (Bezot 1963; Carberry *et al*. 2001). Temperate crops can also be sown at regular time intervals during the favourable sowing windows (Masle *et al*. 1989). Part of the varieties demonstrate a linear, quantitative response to the variable daylength resulting from successive sowing months. Some qualitative (or absolute) photoperiodic varieties experience floral induction inhibition for months, with sequential occurrences grouped together in the same month when daylength fall below a threshold. However, in qualitative short-day sorghum varieties, flowering is initiated again in early July when the days are at their longest. A mobile threshold in a hyperbolic relationship with plant age has been proposed to account for this pattern (Folliard *et al*. 2004; Dingkuhn *et al*. 2008). All of these models are only valid at a single latitude. This is because they predict that flowering time changes proportionally to changes in daylength due to latitude, which contradicts the observed stability of flowering time across latitudes (Curtis 1968; Clerget *et al*. 2021). Building on the dSR/dSS hypothesis and driven by numerous observations that did not align with previous models, our team developed a new model relating three components of the solar photoperiod — daylength (DL), and daily variations in sunrise and sunset times (dSR and dSS) — to the timing of panicle initiation. Panicle initiation is triggered in an embryonic cell mass at the stem apex, meaning the entire process from plant emergence is strictly developmental. In the absence of severe stress and within an adequate temperature range, this process is independent of temperature due to homeostatic adjustments (Clerget *et al*. 2021). The panicle then grows until heading and flowering, and this phase is sensitive to temperature and water supply. The model was calibrated using data from contrasting sorghum and rice varieties sown monthly at various latitudes in the tropics. This model fitted the panicle initiation (PI) data well, particularly when the duration to PI was prolonged for November and December sowings. For the first time, a model accurately fitted observations at latitudes very different from the latitude of the calibration site.

However, for sorghum, the 2021 model did not consider why observations were much more variable for sowings done in Mali from November to February than at any other time of year. Additionally, two sets of model parameters were used to improve the model’s accuracy: one set covering October to June, and the other covering July to September. Nevertheless, this approach was not supported by any biological hypothesis. Lastly, the intuition that the qualitative photoperiodic responses could be set early in the plant life cycle, as the phyllochron does, was explored (McMaster 1997; Clerget *et al*. 2008; Gutjahr *et al*. 2013). This paper aims to propose solutions or hypotheses that address these three questions. Importantly, the dSR/dSS effect should result in different photoperiodic responses in the northern and southern hemispheres. Such a pattern was observed in recent sorghum variety trials conducted in Madagascar and in Argentina, the results of which are reported here.

## MATERIALS AND METHODS

### Data acquisition

#### Variety adaptation trial in Southern Madagascar

The experimental sorghum trial took place at the Centre de Production de Semences d’Agnarafaly (CPSA) in the district of Amboasary (24°55’19’’ S; 46°13’30’’ E; 42 m asl), which is located near the boundary with the Androy region in the humid zone of the Anosy. Average annual rainfall is 1030 mm and average monthly temperatures ranging from 25 °C to 27 °C during the cropping season (Supplementary Fig. S1). The climate is marked by two distinct seasons: a long dry season from April to November and a short rainy season from December to March (*Monographie Région Anosy* 2013). The soil is sandy clay in texture and the previous crop was the leguminous plant lablab bean (*Dolichos lablab*).

Soil preparation involved ploughing, clod breaking, clearing, and levelling, followed by the incorporation of 5 t/ha of compost. Eighty-six early to mid-early sorghum varieties and breeding lines from CIRAD and ICRISAT gene banks and breeding programmes were sown in a non-replicated nursery (Supplementary Table S1). Depending on seed availability, each variety was sown in one to three rows of five meters, with 80 cm between rows and 50 cm between planting hills. Prior to the sowing date on 14 November 2020, a pre-sowing irrigation of 20 mm was applied. Four to six seeds were sown per hill and later thinned to three plants per hole, 15 days after emergence. Due to poor germination and pest damage, some of the plots were resown on 5 December. Only 120 mm of precipitation was recorded between October 2020 and January 2021. In the absence of adequate rainfall, the plots were surface irrigated once per week with approximately 53 mm of water over a 33-minute period per irrigation. The first weeding took place 10–15 days after emergence, with additional weeding and insecticide applied as required. Mineral fertilisation was carried out during the first weeding, involving the application of 100 kg/ha of NPK 9-12-12 fertiliser on 30 November 2020. The dates on which 50% of the plants reached the heading and flowering stages were recorded for each plot and plant height was recorded at harvest on three plants per plot and averaged.

#### Variety adaptation trial in Central Argentina

During the EU-funded project ‘Sweetfuel’, as part of twinning activities between European partners and Argentina, CIRAD shared 70 early to mid-late cultivars with various genetic backgrounds and geographic origins with the sorghum research programme at the National Agricultural Technology Institute (INTA) for testing their potential use in developing new silage and sweet materials. The seeds were sown in Manfredi, Córdoba (31°49’S; 63°45’S) on 7 December 2011 with an experimental planter. The soil is a silty loam Entic Haplustoll (USDA Soil Taxonomy) and the climate is temperate, with a mean annual temperature of 16°C (Fig. S1). The annual average precipitation is 744 mm, predominantly concentrated between October and March. The flowering dates of 38 cultivars were recorded.

#### Monthly sowings in West Africa

Twenty-eight sorghum varieties were monthly sown at the Samanko ICRISAT research station near Bamako, Mali (12°34′ N, 8°04′ W, 330 m asl) from 2000 to 2008 (Supplementary Table S2).

Five series of monthly sowings on approximately the tenth day of each month were conducted from July 2000 to May 2009. Three West African cultivars, ‘CSM 335’, ‘IRAT 174’ and ‘Sariaso 10’, were used in the first series (26 months from Jul 2000 to Aug 2002). Twelve cultivars from West and Central Africa and Madagascar, plus ‘CSM 335’ as a control, were used during a second series (29 months from Nov 2002 to Mar 2005). Seven East African cultivars plus ‘CSM 335’, ‘IRAT 174’ and ‘Sariaso 10’ were used during a third series (24 months from Dec 2006 to Nov 2008). Finally, three durra Muskwaari cultivars from Central Africa (25 months from Oct 2006 to Oct 2008) and three durra cultivars from India and Ethiopia (15 months from Jun 2007 to Aug 2008) were tested.

Individual plots consisted of four 5-m-long rows, sown at 0.75 m × 0.20 m spacing. Split fertilization doses were applied to secure the optimum growth of the plants. During the dry season, irrigation was administered twice weekly, ensuring optimal moisture levels in the soil throughout the study. Hand-weeding and insecticides were used when necessary. The number of leaves that emerged from the whorl on the main culm was recorded weekly from ten specifically labelled plants from two central rows per plot. Two plants per variety were initially sampled every week from seedling emergence to PI, and then at longer intervals for the very late varieties. The plants were dissected in order to count the number of leaves that had been initiated at the apex and to record the PI date. Air temperature and relative humidity at 2 m, soil temperature at 10 cm depth, solar radiation and rainfall were continuously at a weather station located 800 m from the experimental fields.

### Data analysis

#### Aligning comparable values of northern reference and southern recorded heading dates

Northern reference heading durations were recorded in experiments involving sowing dates either in early July in West Africa (2 weeks after the June solstice) or in mid-May in Montpellier or by USDA in Kansas (5 weeks before the June solstice). The Madagascar experiment was sown in mid-November, five weeks before the December solstice — the symmetrical date to the sowings in Montpellier and Kansas — so that the flowering times could be compared. Conversely, the Madagascar experiment was sown seven weeks before the symmetrical date for West African sowings, in mid-January. Longer durations to heading of photoperiod-sensitive varieties are expected in response to this earlier sowing date (Clerget *et al*. 2007). The photoperiod-sensitivity index (Kp) is proportional to the duration to heading recorded on sowings from 9 July, within the range of 49.5 to 98.6 days. Kp was estimated as: Kp = -1.0081 + 0.0204 x min(max(49.5, HD_WCA), 98.6) where HD_WCA is the reference duration to heading for early July sowing in WCA. Based on a 9 July sowing, the duration to heading for a 14 May sowing (56 days earlier) was estimated as:

HD_WCA_14May = HD_WCA + Kp x 56

In Manfredi (Argentina), the sowing dates were two weeks before the solstice (7 December), compared to five weeks in Montpellier (France) (around 15 May). Kp relationships cannot be used because they were not calibrated at these latitudes. Due to the photoperiodic response of the mid-late varieties used, a delay of three weeks delay is expected to decrease the duration to Pi and to flowering time in Montpellier by about 10 days (Clerget *et al*. 2021). To make a fair comparison, these 10 days were subtracted from the flowering time in Montpellier. Conversely, the sowing dates in Manfredi and Saria (Burkina Faso) were similar, so the durations to flowering can be compared.

#### Estimation of the duration from sowing to PI per plant

In all experimental plots conducted in Samanko between 2000 and 2009 panicle initiation date (PI) was recorded weekly by dissecting 1-2sampled plants. The dynamics of leaf emergence over time were linear, bilinear or trilinear, mostly depending on the total number of leaves. The number of leaves observed on each plant was regressed against the elapsed thermal time from emergence using segmented models, with the parameters estimated iteratively. Then, the thermal time at PI (TTPI) was estimated as the thermal time at the appearance of the leaf aLN synchronous with the initiation of the leaf iLN regressed as

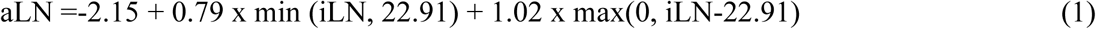

(Supplementary Fig. S2 calibrated on the 28 varieties), with iLN being the rank of the flag leaf of the main stem (Clerget *et al*. 2008). Lastly, the PI date and its duration in days were extracted from the TTPI table.

The same analysis was previously done on the same dataset of 91,827 observations of the number of leaves already appeared per sowing plot. As the record of the upper leaf on the stem ceased after two weeks, once the leaf blade had fully expanded, this method eliminated all early panicle initiation outliers and provided an estimated average date for the later initiations in the 761 observed plots. This resulting dataset was used with other datasets in Clerget *et al*. (2021). In this paper, the estimated initiation date of each of the 6,806 observed plants was computed, thus keeping interplant variability in each plot.

### Statistical analysis

For the monthly sowings in Bamako, the normality of the data distribution was tested prior to Glm ANOVAs, using SAS 9.4. RStudio 4.5.2 was used to compute t-tests for differences in flowering dates between the northern reference dates and the data from Madagascar and Argentina, and the research model. The confidence interval for the mean difference in flowering date was used to create a confidence area around the 1:1 slope in graphs comparing flowering dates in the northern and southern hemispheres. The root mean square error (RMSE) was used to compare the model outputs.

### A research model integrating four potential photoperiodic responses, including qualitative responses set at an early stage

A research model was developed to test the accuracy of the hypothesis that different photoperiodic responses coexist within each cultivar. It postulates that there were two observable quantitative and two observable qualitative photoperiodic responses throughout the year.

The effect of temperature was not a major factor in determining the PI date in these experiments due to the quite stable mean temperature in Samanko (25.9 ± 4.1 °C, Supplementary Fig. S3) and the observed adaptation of the plants to temperature during the early stages (Vergara and Chang 1985; Birch *et al*. 1998; Hu *et al*. 2026). Consequently, this study, which focused on plant development until panicle initiation, did not consider the effects of temperature. Conversely, temperature significantly impacts the length of the subsequent growth phase, from panicle initiation to flowering (Ritchie and NeSmith 1991). The two quantitative responses (Kt1 and Kt2) were computed as according to Summerfield *et al*. (1992) but with additional factors. Initially, the expected duration of the the photoperiod-sensitive phase (PSPi) under the photoperiod of the day i (PPi) was a function f(PPi) of three components of the daily photoperiod (the daylength, DL) and the changes in sunrise time (dSR) and sunset time (dSS)). The quantitative responses over the year were then,

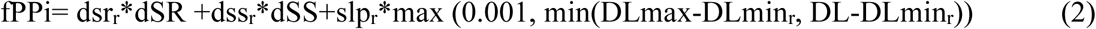

where dSR and dSS are the change in sunrise and sunset times of the day, DL is the daylength (between sunrise and sunset), DLmin_r_ and DLmax are the thresholds of the variety reaction to DL, and slp_r_, dsr_r_ and dss_r_ are the sensitive coefficients to each of the three PP components. After the end of the photoperiod insensitive phase (PIP), plants accumulate a daily progress (dPI) towards PI, according to the equation dPIi=1/f(PPi). PI occurrs when the sum of dPI reaches 1.

In addition, Kt1 was calibrated over two annual periods, perceived at the end of the PIP, as done in the Photoperiodism 2.0 model (Clerget *et al*. 2021). The two periods begin and end on the equinoxes. Daylight hours exceed 12 hours during period 1 and fall below 12 hours during period 2. Thus, two sets of each parameter were created for Kt1, except for PIP and DLmax. QL1 was only observed during period 1, and Kt2, QL2 and QL2s were only observed during period 2. All variety parameters were recurrently adjusted to fit the observed data.

Finally, an attempt was made to remove DL from the potentially overparametred relationship. Consequently, PSPi became a linear function of daily changes in sunrise and sunset times (dSR and dSS), as well as a constant (Cstp), as presented below:

1. Quantitative response 1 (Kt11 and Kt12) during periods 1 and 2:

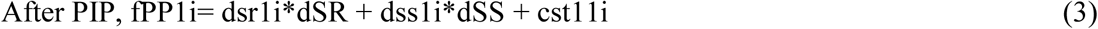 Where and dsr11, dss11, cst11, dsr12, dss12 and cst112 are the sensitive coefficients to each of the two PP components and the constants, during periods 1 and 2.
2. Quantitative response 2 (Kt2) Conversely to Kt1, this second quantitative response was only observed during period2 from the autumn equinox to the spring equinox.

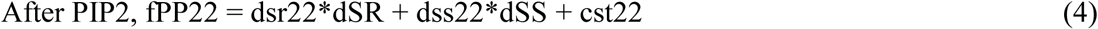 where dsr22, dss22 and cst22 are the sensitive coefficients to each of the two PP components and the constant during period 2. Two coincidence qualitative responses based on waiting for a specific value of dSR+dSS (Beta) set at the end of the photoperiod insensitive phase (PIP) in response to dSR and dSS. The dSR+dSS values can indifferently increase or decrease when Beta is reached and PI is triggered.
3. Qualitative response 1 (QL1) in January to June sowings The application period spans from two days after the winter solstice (as dSSR peaks one day after the solstice) to the summer solstice at the end of PIP stage (pip), when dSRpip < dSSpip and dSSRpip < 61.55.

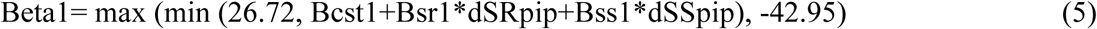 where dSRpip, and dSSpip are the values of dSR and dSS at the end of PIP, 26.72 is 0.08 lower than the maximum value of dSR+dSS on the summer solstice, is 0.02 higher than the minimum value of dSR+dSS on the spring equinox, and Bcst1, Bsr1 and Bss1 are the three parameters of the linear relationship between dSR+dSS at PI and dSRpip and dSSpip. In mid-late varieties, when dSR+dSS at PIP is decreasing, PI is reached before the summer solstice when dSR<dSS and dSR+dSS is increasing and dSR+dSS>Beta; reversely when dSR+dSS at PIP is increasing, PI is reached after the summer solstice when dSR>dSS and dSR+dSS is decreasing and dSR+dSS<Beta. In late varieties, PI is always reached after the summer solstice when dSR>dSS and dSR+dSS is decreasing and dSR+dSS<Beta. Parameters of the two possible models are optimised for each variety, and the one with the lower root mean square error (RMSE) is kept. In some varieties, PI marked QL1b occurred on an alternative date for February and March sowings but at the same Beta value, thus before the summer solstice, while PI noted QL1 occurred after the solstice. Beta1 parameters were recurrently adjusted for each variety on observed PI data from January until June emergence dates. However, as dSRpip and dSSpip increased with latitude, so did Beta1, resulting in an increase in the predicted dSR+dSS at PI. This led to an earlier prediction of the time of PI, which did not align with the observed times of PI. Indeed, the observed times were largely independent of latitude. This undesirable effect was corrected by adding a daily variable linear function to Beta1: dl1*DLday, where dl1 is a parameter and DLday is the day length. Some data recorded in Montpellier enabled all parameters, including dl1, to be readjusted to fit data at two latitudes in two cultivars: CSM 335 and Souroukoukou.
4. Qualitative response 2 (QL2) in November to February sowings The application period is centered on the winter solstice and spans from autumn equinox (the first day when dSR > dSS and dSR+dSS is increasing) to the spring equinox (the last day when dSR < dSS and dSR+dSS is decreasing) at the end of PIP stage (PIP),

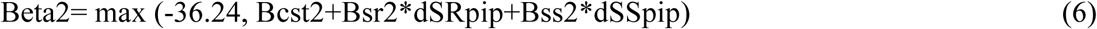 where dSRpip, dSSpip, dSR+dSSpip, are the values of the three PP factors at the end of PIP, - 36.24 is 0.06 higher than the minimum value of dSR+dSS on the spring equinox, and Bcst2, Bsr2 and Bss2 are the three parameters of the linear relationship between dSR+dSS at PI and dSRpip and dSSpip. PI was reached when dSR<dSS and dSR+dSS decreases and dSR+dSS<=Beta2.
5. Qualitative response 2 in the Tanzanian southern varieties (QL2s) in November to February sowings. The period of application is similar to that of QL2.

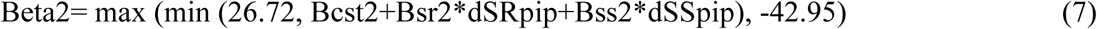 where dSRpip, dSSpip, dSR+dSSpip, are the values of the three PP factors at the end of PIP, 26.72 is 0.09 lower than the maximum value of dSR+dSS on the summer solstice, -42.95 is 0.018 higher than the minimum value of dSR+dSS on the autumn equinox, and Bcst2, Bsr2 and Bss2 are the three parameters of the linear relationship between dSR+dSS at PI and dSRpip and dSSpip. When dSR+dSS at pip is increasing, PI is reached before the summer solstice when dSR<dSS and dSR+dSS is increasing and dSR+dSS>Beta; reversely when dSR+dSS at pip is decreasing, PI is reached after the summer solstice when dSR>dSS and dSR+dSS is decreasing and dSR+dSS<Beta.

Plant PI observations within each variety x monthly sowing date combination were allocated to one of the six photoperiodic response categories if their distribution significantly diverged from normality and they constituted significantly different groups. The number of responses was based on the number of strictly distinct means obtained from the post hoc SNK test. Otherwise, all data were allocated to a single response. Allocation to a specific response was done manually.

For comparison purposes, the previous Photoperiodism 2.0 model was re-computed using the 6806 plant PI instead of the initial by-plot PI.

## RESULTS

### Delayed heading of cultivars transferred from the Northern to the Southern Hemisphere Early cultivars transferred to Madagascar headed very late

The duration to heading of 74 early- to mid-early varieties and lines developed in the Northern Hemisphere and three cultivars from Equatorial/Southern Africa was recorded in an adaptive trial carried out in the Anosy region, southern Madagascar. The plants grew well during the vegetative phase, reaching heights of 3 to 3.5 metres for the late varieties (Table S1). Irrigation was stopped 140 days after sowing, by which time the early and mid-early varieties had already been harvested. Twelve cultivars had not yet headed.

Thirty-one accessions exhibited similar heading durations in Madagascar than in the northern hemisphere, while 33 accessions headed significantly later than the northern reference date and 13 early accessions headed significantly earlier than the northern reference (Fig. 2). The mean observed difference in heading times between the northern and southern hemispheres was 12.708 days (P < 10^-4^), with asymmetric distribution, and the 95% confidence interval bootstrap for the null difference was [-6.7, 7.2]. The heading time of three cultivars from Equatorial and Southern Africa in Madagascar remained close to the Northern reference (squares, Fig. 2).

**Fig 1:**
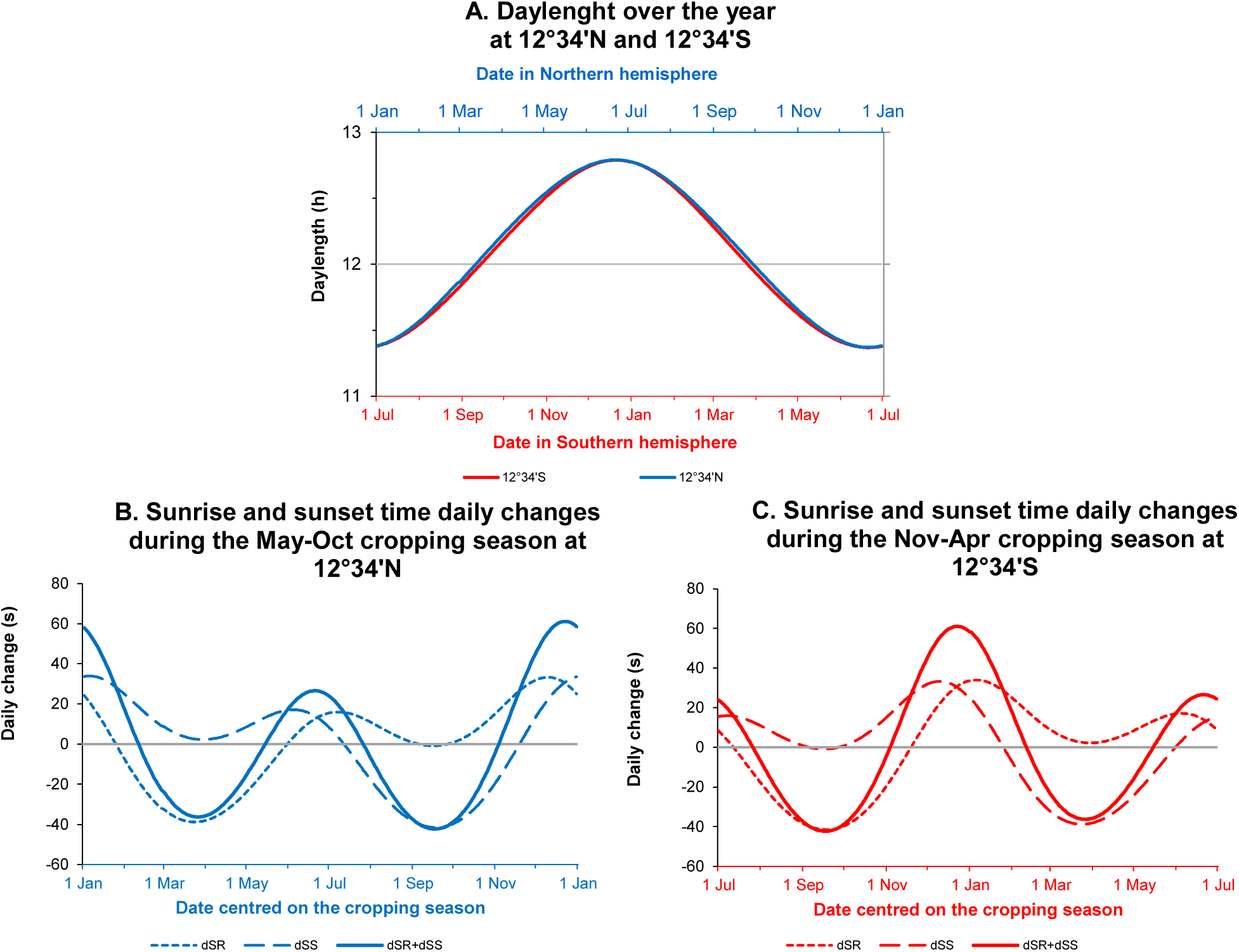
Daily values of four photoperiod components-Daylength (A), and daily changes of sunrise (SR) and sunset (dSS) times and of their sum (B and C)-at two places located at symmetrical latitudes in the Northern hemisphere (Bamako, 12°34’N) and in the Southern hemisphere (hypothetic location, 12°34’S).

**Fig. 2:**
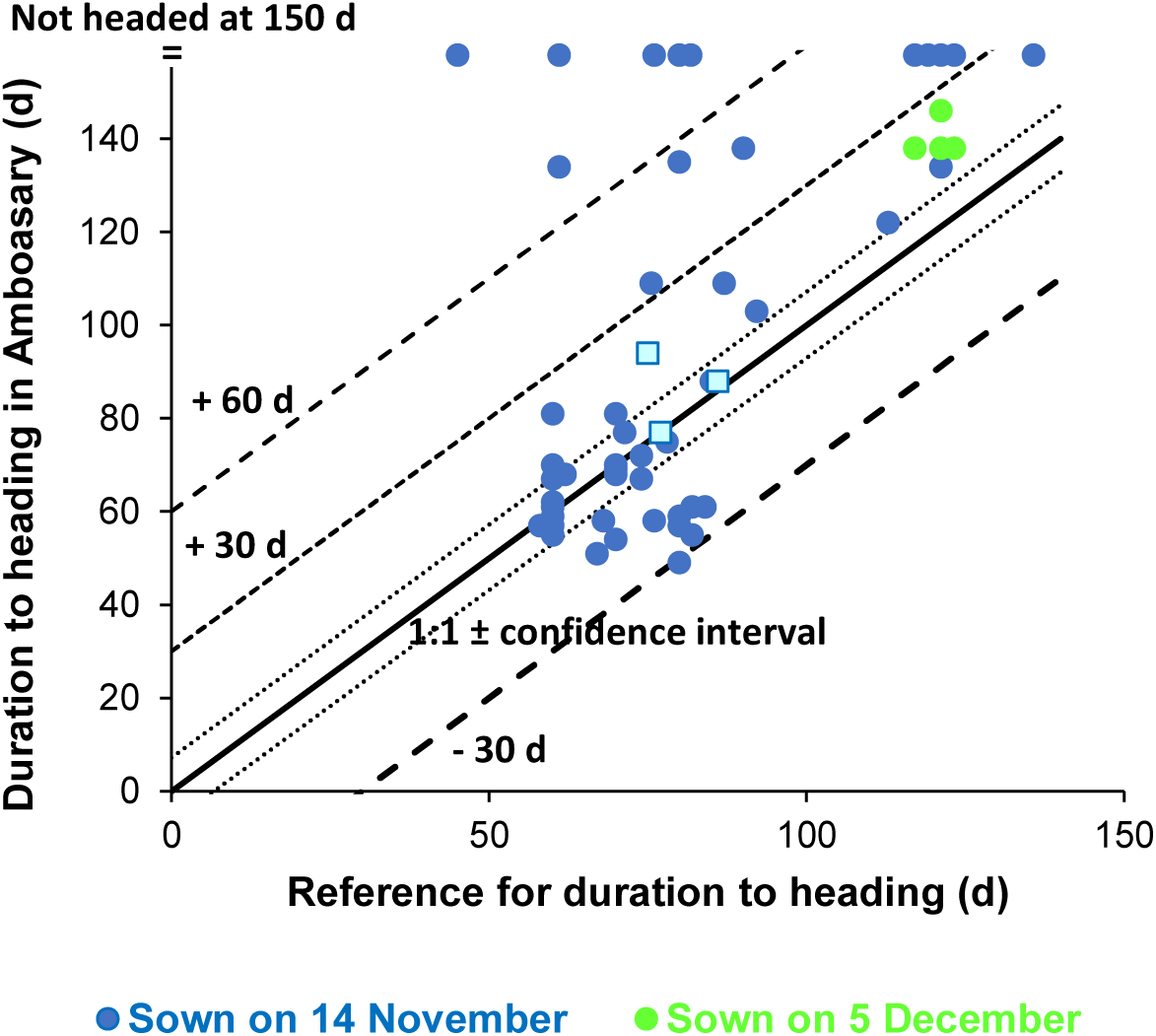
Comparisons between the reference heading date of 74 early sorghum cultivars from the Northern Hemisphere (blue and green dot) and 3 cultivars from the Equatorial or Southern Africa (square) when sown in May in the Northern Hemisphere and their heading date when sown in Nov-Dec in southern Madagascar (24°55’ S).

#### Flowering time of many cultivars was significantly delayed in Argentina

The duration to flowering of 29 mid-early cultivars cropped either in the northern hemisphere or in Manfredi, Argentina could be compared. All but three of the observed flowering times were later in Manfredi than in Montpellier or Saria (Supplementary Fig. S4 and Supplementary Table S3). The mean observed difference in heading times between the northern and southern hemispheres was 17.97 days (P < 10^-8^), and the 95% confidence interval for the null difference was ± 4.153 days. The cultivars from Botswana, Lesotho, and Eswatini also flowered later in Manfredi than in the northern hemisphere.

### Alternative dates of panicle initiation in November to February sowings in the CSM 335 variety

The Malian guinea landrace CSM 335 was used as an experimental check, with its phenology recorded over a period of just over six complete years (76 months) across three distinct series of consecutive sowing months. For sowings from March to October, the duration to PI was highly repeatable over six experimental years with standard deviations comprised between 5 and 11 days (Fig. 3A&B and table 1). Each monthly set of observations was normally distributed, except in June and October. However, year always had a significant, albeit moderate, effect.

**Fig. 3:**
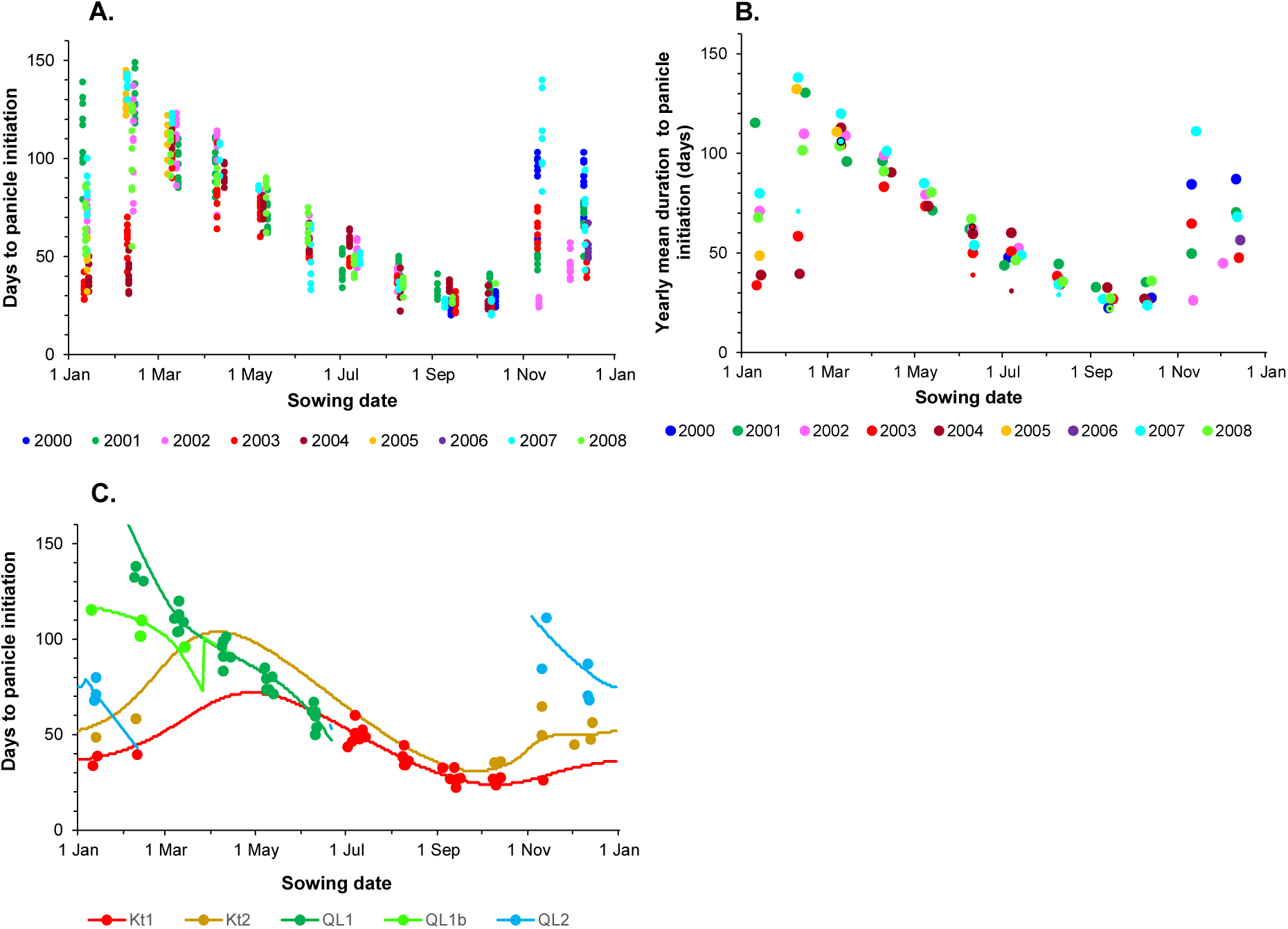
Distributions by sowing date in CSM 355 (A) of 661 per plant values for durations to PI recorded in monthly sowing series during six to seven years, (B) of their 76 per plot monthly mean durations to PI and seven extreme outliers (small dots), and (C) allocation of the 76 mean durations to PI to four photoperiodic response types: quantitative (Kt1 and Kt2) and qualitative (QL1 and QL2). Curves show the daily predicted durations to PI of the four response types.

**Table 1:**
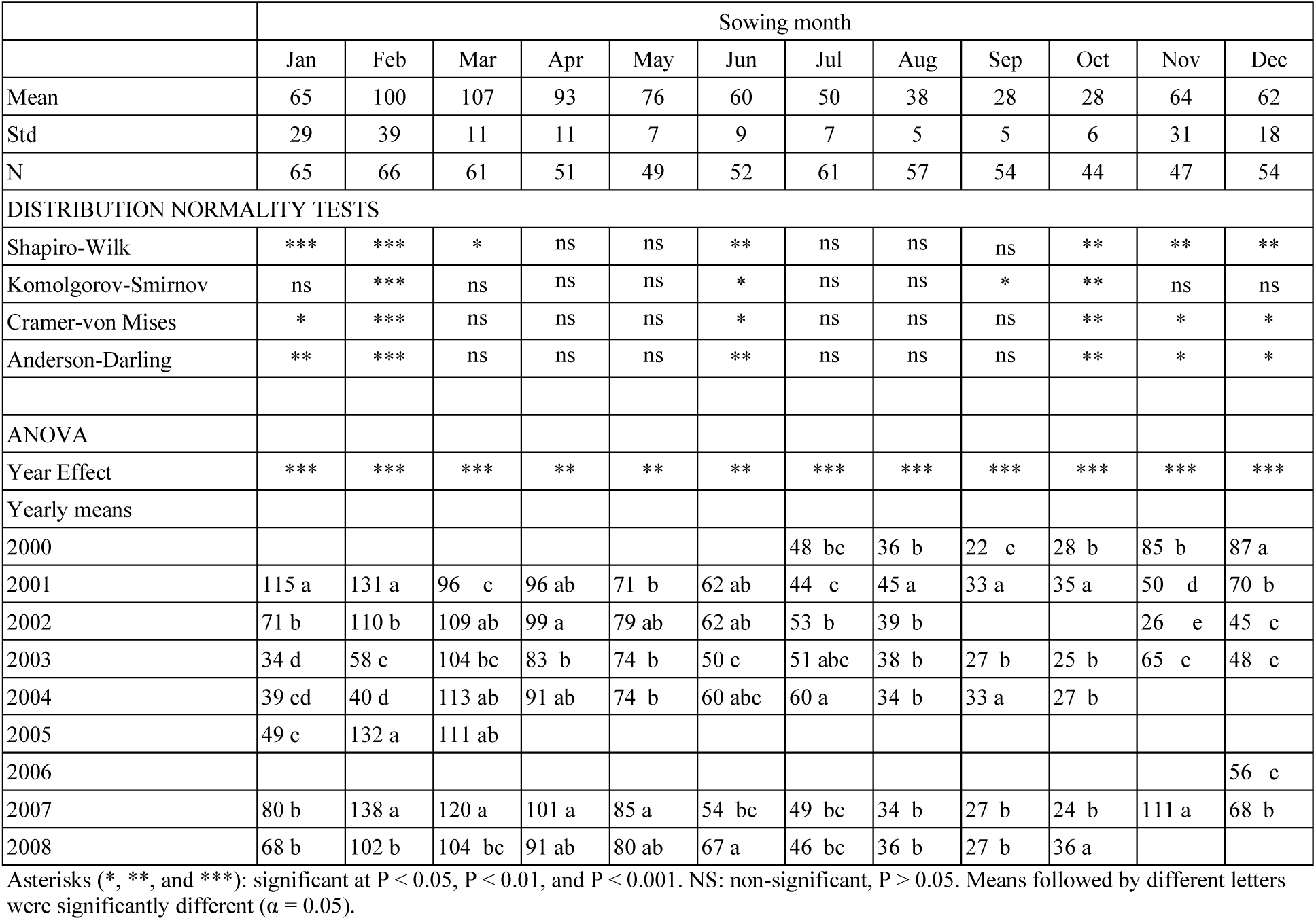
Monthly data distribution with normality tests, ANOVA of the year effect on the duration to panicle initiation in CSM 335, and yearly mean durations sorted with SNK test.

|  | Sowing month |  |  |  |  |  |  |  |  |  |  |
|---|---|---|---|---|---|---|---|---|---|---|---|---|
|  | Jan | Feb | Mar | Apr | May | Jun | Jul | Aug | Sep | Oct | Nov | Dec |
| Mean | 65 | 100 | 107 | 93 | 76 | 60 | 50 | 38 | 28 | 28 | 64 | 62 |
| Std | 29 | 39 | 11 | 11 | 7 | 9 | 7 | 5 | 5 | 6 | 31 | 18 |
| N | 65 | 66 | 61 | 51 | 49 | 52 | 61 | 57 | 54 | 44 | 47 | 54 |
| DISTRIBUTION NORMALITY TESTS |  |  |  |  |  |  |  |  |  |  |  |  |
| Shapiro-Wilk | *** | *** | * | ns | ns | ** | ns | ns | ns | ** | ** | ** |
| Komolgorov-Smirnov | ns | *** | ns | ns | ns | * | ns | ns | * | ** | ns | ns |
| Cramer-von Mises | * | *** | ns | ns | ns | * | ns | ns | ns | ** | * | * |
| Anderson-Darling | ** | *** | ns | ns | ns | ** | ns | ns | ns | ** | * | * |
| ANOVA |  |  |  |  |  |  |  |  |  |  |  |  |
| Year Effect | *** | *** | *** | ** | ** | ** | *** | *** | *** | *** | *** | *** |
| Yearly means |  |  |  |  |  |  |  |  |  |  |  |  |
| 2000 |  |  |  |  |  |  | 48 bc | 36 b | 22 c | 28 b | 85 b | 87 a |
| 2001 | 115 a | 131 a | 96 c | 96 ab | 71 b | 62 ab | 44 c | 45 a | 33 a | 35 a | 50 d | 70 b |
| 2002 | 71 b | 110 b | 109 ab | 99 a | 79 ab | 62 ab | 53 b | 39 b |  |  | 26 e | 45 c |
| 2003 | 34 d | 58 c | 104 bc | 83 b | 74 b | 50 c | 51 abc | 38 b | 27 b | 25 b | 65 c | 48 c |
| 2004 | 39 cd | 40 d | 113 ab | 91 ab | 74 b | 60 abc | 60 a | 34 b | 33 a | 27 b |  |  |
| 2005 | 49 c | 132 a | 111 ab |  |  |  |  |  |  |  |  |  |
| 2006 |  |  |  |  |  |  |  |  |  |  |  | 56 c |
| 2007 | 80 b | 138 a | 120 a | 101 a | 85 a | 54 bc | 49 bc | 34 b | 27 b | 24 b | 111 a | 68 b |
| 2008 | 68 b | 102 b | 104 bc | 91 ab | 80 ab | 67 a | 46 bc | 36 b | 27 b | 36 a |  |  |

Asterisks (\*, \*\*, and \*\*\*): significant at P < 0.05, P < 0.01, and P < 0.001. NS: non-significant, P > 0.05. Means followed by different letters were significantly different (α = 0.05).

In contrast, interannual variability was much higher for sowings from November to February, with standard deviations of 18 to 39 days, and non-normality of the dataset distribution. The analysis of individual plant durations to PI showed a significant year effect on this variability and the distribution of the yearly mean durations into four, four, five, and three totally distinct groups of annual means in January, February, November and December sowings, respectively (Table 1). While no group was entirely distinct from the others for the March to July sowings, two groups were fully distinct for the August to October sowings. The distribution of the means also diverged from normality in October. It was consequently hypothesised that the contrasting groups of mean durations to PI resulted from four types of photoperiodic response: quantitative (Kt1, over the year, and Kt2 in Nov-Feb), and qualitative (QL1 in Jan-Jun, and QL2 in Nov-Feb), with QL1b being an early alternative to QL1 (Fig. 3C). In general, only the more photoperiod-sensitive response of the ongoing season was expressed in Mar-Sept. Conversely, two to three significantly different durations to PI were found for sowings from November to February in a further 22 varieties (Supplementary Table S4).

Consequently, up to four photoperiodic responses were hypothesised for a further 27 varieties, with QL2s being a variation of QL2 in five varieties (Fig. 4).

**Fig. 4:**
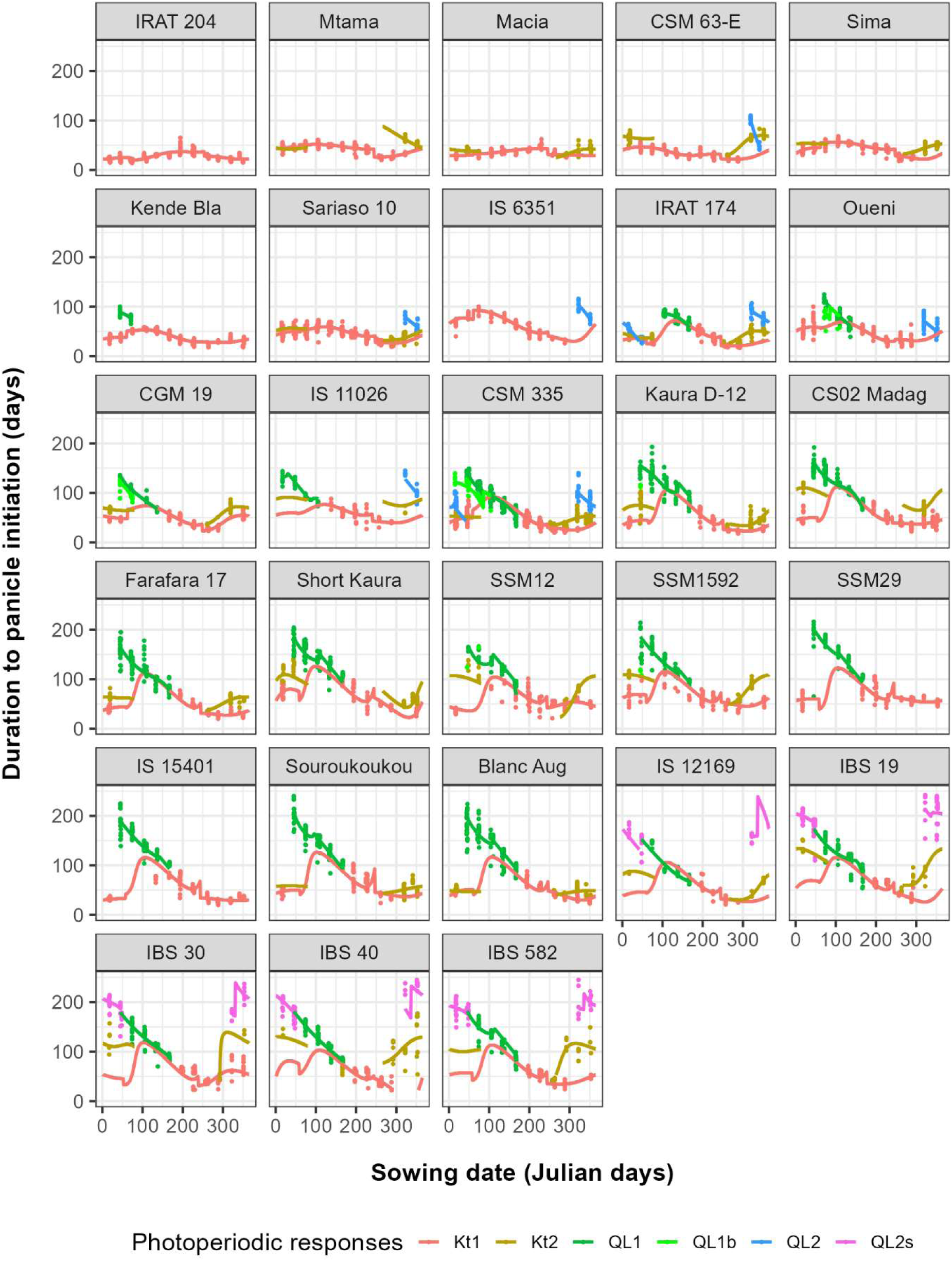
Display of the 6,806 durations to panicle initiation (PI) observed in 28 sorghum varieties sown monthly in Bamako, Mali, from 2000 to 2008, alongside their four potential photoperiodic responses: two quantitative (Kt) and two qualitative (QL). The models used to predict the four daily responses only considered the dSR and dSS photoperiodic variables. QL1b is an alternative duration to QL1 when PI is triggered before rather than after the summer solstice. QL2s is a very late QL2 response that is specific to a group of five varieties

Additionally, one outlying plant sown in February 2007 had a much shorter duration to PI than all the other plants but close to mean duration in the February 2003 sowing (Fig. 3B). Similarly, an earlier outlier plant was observed in IBS 30 in the May sowing, and in SSM 29 in the February sowing (Fig. 4). These rare but recurring events demonstrated that few plants evaded the yearly response and switched to a different one. Therefore, plants of the same variety exhibited up to four durations of PI in response to the photoperiod, which primarily depended on the year of sowing and, to a lesser extent, on the individual plants.

### Alteration of the light spectrum by the Harmattan wind affected the perception of dSR and dSS

The interannual variability from November to February was primarily observed in the second series of sowings across the three winter seasons (2002–03, 2003–04, and 2004–05). The mean duration to PI for this season was significantly lower than that of the other seasons (P < 0.001). This called into question the possibility of a change in cultivar used. Actually, none of the cultivars in this series showed a long duration of PI for sowings made between November and February, despite the existence of seven cultivars that were later than CSM 335 for February to June sowings. Unlike the other experimental years, 2003–2006 saw frequent dusty Harmattan winds in Bamako, with mean temperatures above seasonal averages (23.8 °C compared to 22.4 °C in other years in Samanko) (Fig. 5). The Harmattan is a northeasterly wind that blows from the Sahara across West Africa from December to March. However, it has been demonstrated that changes in the light spectrum have a significant impact on the time to PI of the CSM 335 sorghum variety when sown from December to February and grown either in the field or in a greenhouse at similar temperatures (Clerget *et al*. 2012). Indeed, the glass in the greenhouse used in the study filtered out blue and UV waves, which are perceived by phytochromes involved in the flowering pathway (Anuforom *et al*. 2007). Therefore, the dark yellow light during the Harmattan periods, coupled with the accumulation of dust on plant leaves, were likely more responsible for the shorter PI durations observed from 2003 to 2006 than the slight temperature increase. Thus, the shorter durations to PI during the November to February sowings of CSM 335 from November 2002 to February 2005 were probably caused by a recognised climatic difference. Any doubt regarding a possible seed difference can therefore be dismissed. Additionally, besides these three specific years, the interannual variability of the duration to PI remained high during the other years.

**Fig. 5:**
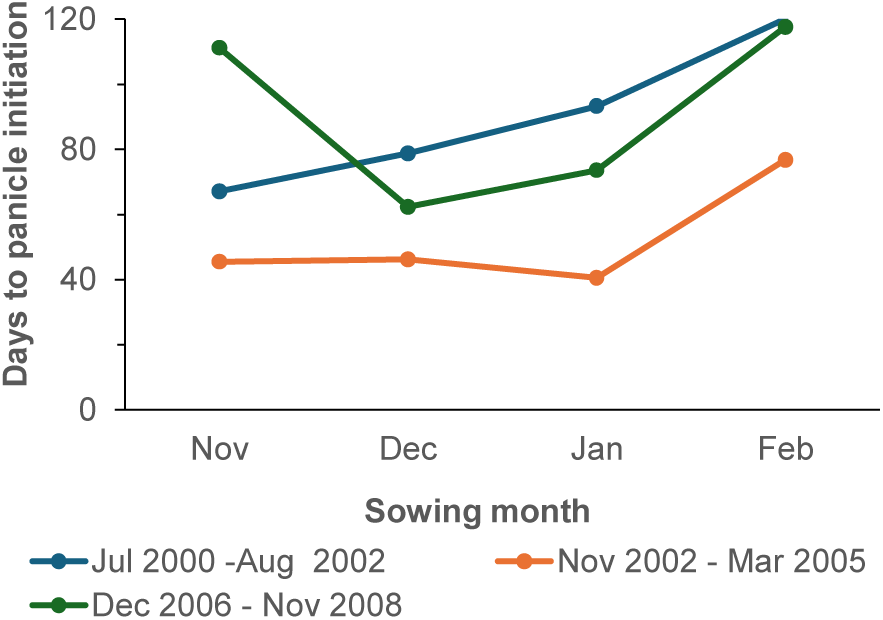
Mean durations to panicle initiation of CSM 335 for November to February sowings, by sowing series.

### Two wide, empty time windows in the yearly distribution of the panicle initiation dates

#### Main photoperiodic response for sowings from February to August

Varieties were sorted by increasing delays to PI for the February to June sowings, and yearly panicle initiation mean dates were plotted per variety and sowing month (Fig. 6A). The start of the timeline of the graph showing the date of panicle initiation has been set to September sowing, when the time to PI is at its shortest for all varieties. Panicle initiation occurred about one month after sowing for the non-photoperiodic variety IRAT 204. Panicle initiation of the other 27 varieties occurred later than in IRAT 204, leading to group them into four photoperiodic classes: 1) In six quantitative photoperiodic varieties the panicle initiation of the February to June sowings was up to 45 days later than in IRAT 204; 2) In eight qualitative photoperiodic varieties, PI of plants sown from March (or April in IRAT 174) to June was inhibited until the summer solstice (21 June) when the sum of sunset and sunrise changes (dSR+dSS) was maximum (Fig. 6C); 3) In nine qualitative varieties, PI of plants sown from February to June was inhibited from the spring equinox (21 March) until 20 July when dSR+dSS was zero, or later. All delayed PI distribution took place between 20 July and the autumn equinox (21 September), when dSR+dSS was at the minimum. 4) In the four Tanzanian varieties, PI of plants sown from January to June was inhibited until 20 July but occurred before 1^st^ September. PI of the November and December sowings could be delayed by up to 7 months while occurring from May to July.

**Fig. 6:**
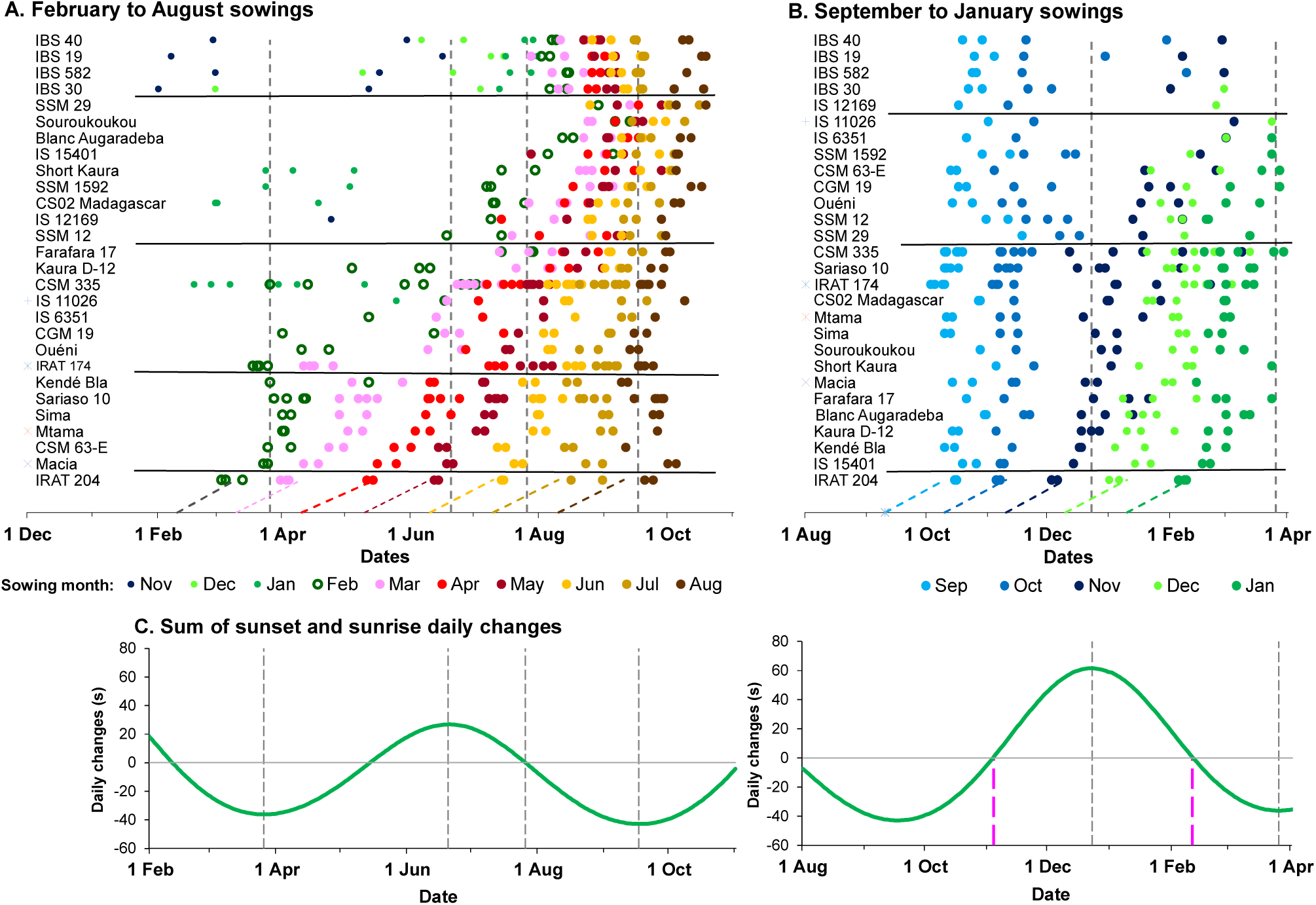
**A**. Distribution of panicle initiation dates of 28 sorghum varieties monthly sown from September to January and sorted by increasing photoperiod sensitivity during this period. **B**. Distribution of panicle initiation dates of 28 sorghum varieties monthly sown from February to August and sorted again by increasing photoperiod sensitivity during this period. Dashed diagonals show one month from the sowing date on the 10^th^ of each month. **C.** Sum of the daily change of the sunset and sunrise times (dSR+dSS) over the year. Vertical grey lines show the date when dSR+dSS values are maximum, zero, or minimum.

#### Secondary photoperiodic response for sowings from November to February

The 28 varieties were sorted again by increasing delays to PI in the November-January sowing (Fig. 6B), which resulted in windows with low PI occurrences leading to group the varieties into three classes: 1) In 14 winter-quantitative photoperiodic varieties the panicle initiation of the November to January sowings was 15 days to 1 month later than in IRAT 204, occurring after the dSR+dSS maximum on 21 December (Fig. 6C). 2) In eight winter-qualitative photoperiodic varieties, the panicle initiation of the November to January sowings was 15 days to 2 months later than in IRAT 204, and the panicle initiation of the October sowing was also delayed in some varieties. With few exceptions, PI of November to January sowings occurred before 21 March when dSR+dSS was minimal. 3) In four Tanzanian and one Ethiopian varieties, PI was inhibited during up to 3 months in October sowings, and up to 8 months in November to January sowings.

#### Relationship between the two seasonal variety photoperiodic responses

The mean durations to panicle initiation for the 28 varieties for the May-June and November-December sowings (before the spring and winter solstices, respectively) were not significantly correlated (ρ = 0.29, P = 0.13). However, the varieties were segregated into four distinct clusters containing thirteen, ten, four and one varieties, respectively (Fig. 7). Durations to PI in May–June and November–December were significantly correlated in the group including IRAT 204 (in blue, ρ = 0.94, P < 0.0001) and not correlated in the other groups (green, ρ = 0.28, P = 0.44; red, ρ = 0.51, P = 0.49). The variety CSM 63E showed a unique pattern. Seven varieties (with names other than SSM) out of the 10 varieties of the group of late varieties for the May-June sowings and early for the Nov-Dec sowings (in green) were tested from Dec 2002 to Mar 2005. During this period, CMS 335 exhibited early panicle initiation from November to February, which was probably caused by the Harmattan winds (see previous paragraph). Like CSM 335, these seven varieties may have exhibited a longer duration to PI and align with the blue group if tested in other sowing series. Conversely, the three ‘SSM’ varieties were tested from Dec 2006 to Nov 2008 and were early for November-December sowings, whereas CSM 335 was late.

**Fig. 7:**
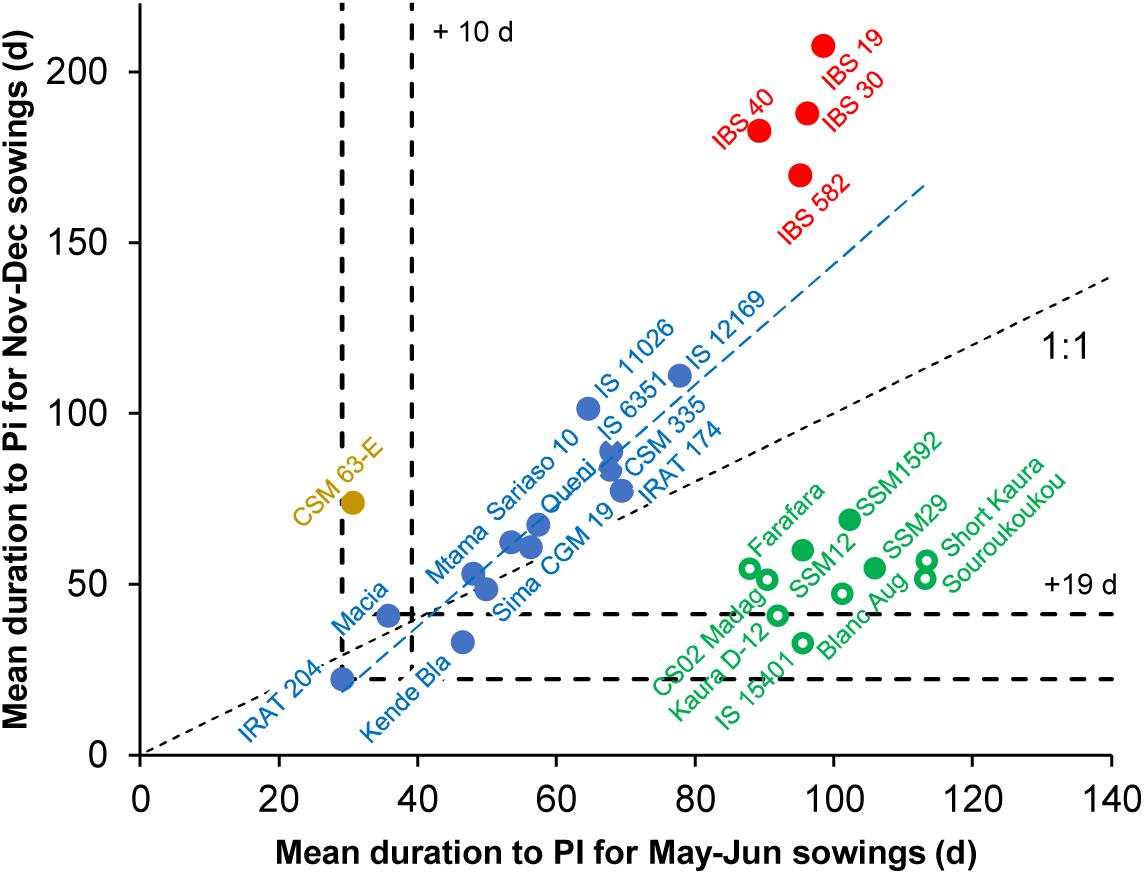
The 28 varieties clustered into four groups based on their mean duration to PI in May-June sowings versus Nov-Dec sowings. Black Dashed lines show the two mean durations to PI of IRAT 204 and their confidence interval (Bonferini post hoc test, α = 0.05). Blue dashed line shows the regression between values from the blue group of varieties.

#### Linear relationship between dSR and dSS at the end of the photoperiod-insensitive phase and dSR+dSS at panicle initiation in qualitative photoperiodic responses

Panicle initiation of highly photoperiod-sensitive varieties sown from January to June were long delayed and then occurred in the same sequence than at sowing during the three months between the summer solstice and the autumn equinox when dSR+dSS continuously decreased (Fig. 6A). Functions between the photoperiod components at the beginning of the photoperiod-sensitive phase and the universally valued dSR+dSS at PI were searched as a support to an eventual coincidence phasic model for this qualitative response. The linear combination Beta = Bcst + Bsr *dSR + Bss * dSS where Beta is the predicted value of dSR+dSS at PI and Bcst, Bsr, and Bss are three optimised parameters fitted the observed dSR+dSS value for CSM 335 well (r Pearson=0.846, P<10^-15^) (Fig. 8).

**Fig. 8:**
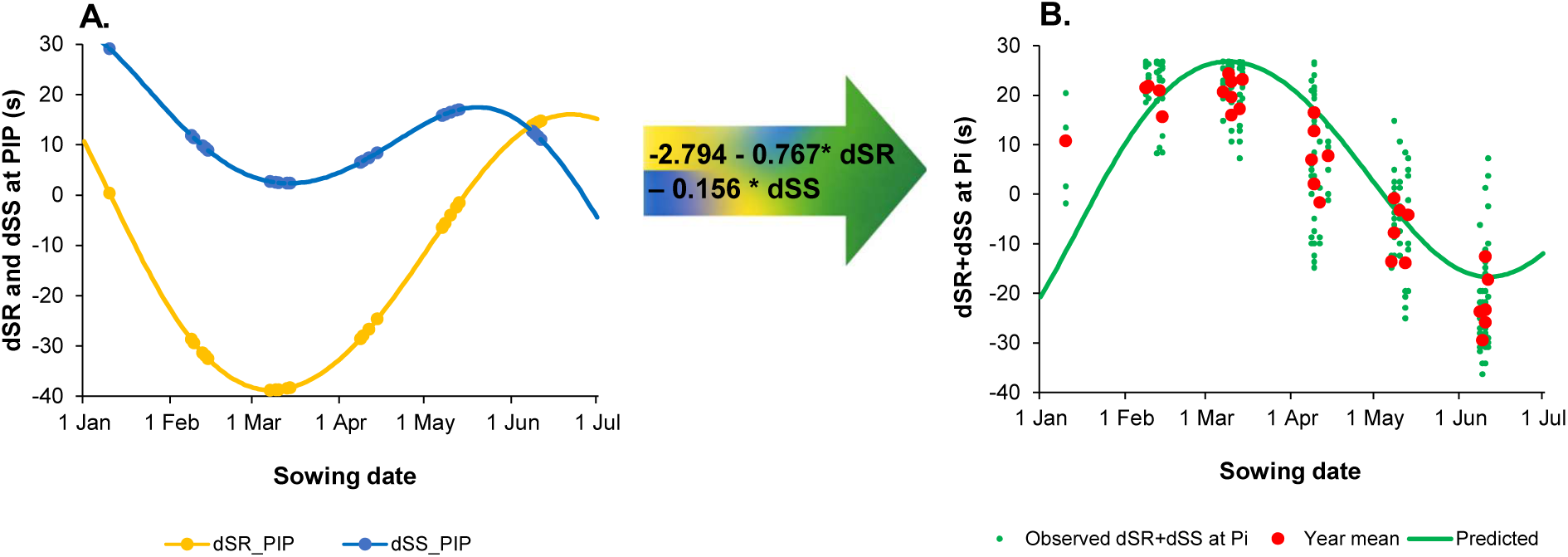
Linear combination of dSR and dSS at the end of the PIP (A) was a good predictor of the observed dSR+dSS at PI (B) of plants having shown a qualitative response to the photoperiod in the variety CSM 335 monthly sown from January to June.

The same linear relationship was applied to November–February sowings of varieties exhibiting a qualitative photoperiodic response (QL2) during this period. The equation was incorporated into the QL1 and QL2 models, with its parameters being optimised alongside all the others.

#### Application of a phenological research model with four potential photoperiodic responses to 28 varieties

Two phenological models, which were based on the results obtained from CSM 335, were calibrated for each of the other 27 varieties using the data recorded in Samanko, Mali (Fig. 6, Supplementary Tables S5 and S6). Two to four photoperiodic responses were manually detected in this panel. Six varieties exhibited a single quantitative response. The other 22 varieties were better fitted with two quantitative responses. The RMSE calculated using the model with only dSR and dSS (noDL: 11.37) was slightly higher than the RMSE calculated using the model with daylength included in the Kt1 and Kt2 equations (DL: 10.77) for 28 varieties (n=6806). The errors of both models were not normally distributed. The distributions of the errors were, however, not significantly different (Wilcoxon paired test: W = 2,291,527; p = 0.2838). Therefore, as both models are equally accurate, the model using DL is overparametrized and the DL factor can be disregarded. The global RMSE of the Photoperiodism Model 2.0 for durations to PI applied to the same dataset was 17.59, in comparison.

The model was then tentatively run at a southern latitude inverse to that at Samanko (for example, in northern Mozambique or Madagascar). This attempt showed that, for sowings during the rainy season (November to January), the mid-early variety Sariaso 10 would have a shorter duration to PI in the Southern Hemisphere than in the Northern Hemisphere (see Fig. 9A). As predicted by QL1, the durations to PI could align with May–June sowings in the Northern Hemisphere for the mid-late CSM 335 variety (Fig. 9B). Conversely, the durations to PI of the late Souroukoukou variety would be considerably longer than in the Northern Hemisphere (Fig. 9C).

**Fig. 9:**
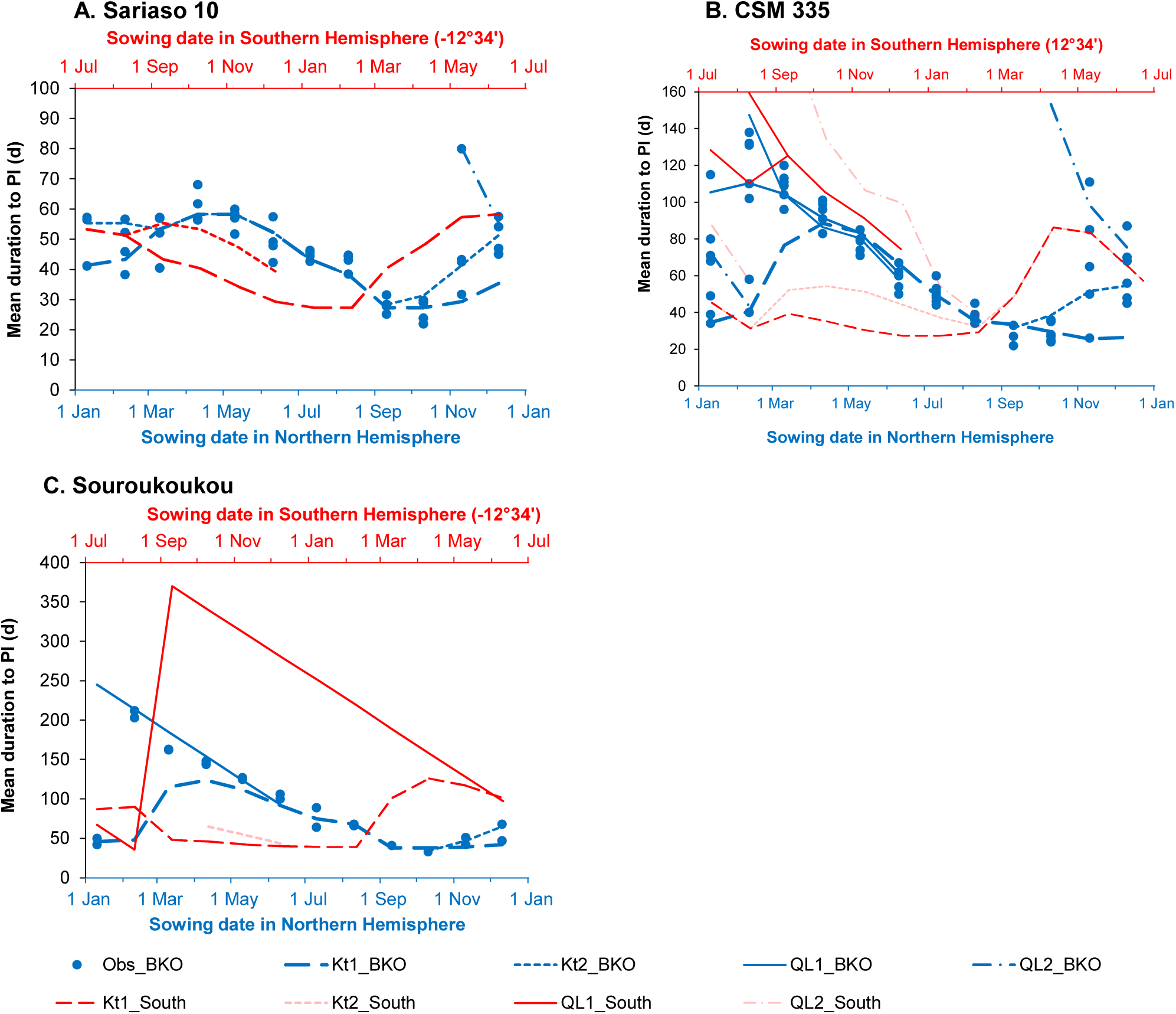
Distributions by sowing date of the monthly means for durations to PI, as observed for three varieties in tropical and temperate locations in the Northern Hemisphere, plotted alongside the daily durations to PI predicted in Bamako (dark blue) and, in a location in the Southern Hemisphere at - 12°34’ (red).

Moreover, after recalibration using data recorded at two latitudes (Samanko and Montpellier), for the two cultivars, the no DL Kt1 model fitted the entire datasets well. (At two latitudes for CSM 335 and Sariaso 10, and at four latitudes for Souroukoukou (Clerget *et al*. 2021). (Data not shown). Conversely, the NoDL QL1 qualitative model, which was calibrated using Samanko data, was not continuously defined at the Montpellier’s latitude. It predicted values for the expected dSR+dSS that were too high, resulting in a PI date that was too early, particularly for sowings in April and May. This undesirable effect could be eliminated by assuming a linear adjustment of the expected dSR+dSS value proportional to the daylength at the time of PI (Supplementary Table S7).

At southern latitudes, the highly photoperiodic cultivar Souroukoukou would experience PI inhibition from September onwards, until the next minimum dSSR value is reached on 17 September. This pattern is somewhat similar to that observed with the four Tanzanian varieties when grown in Samanko, though PI inhibition only began with November sowings, with PI ability recovering between May and July for sowings from November to January. Thus, the exceptionally long durations to PI observed for sowings from November to January in Samanko for these four varieties may have been caused by their inability to adapt to the northern latitude. Information on the PI duration of sowings from May to July in their place of origin would provide an answer, but unfortunately this information is unavailable.

## DISCUSSION

### The effect of the Earth’s hemisphere on floral induction in sorghum

Daylength variation is mainly caused by the earth tilt over its orbital plane. They consequently have symmetrical seasonal values at opposite northern and southern latitudes (Fig. 1A). This is why seasonal photoperiodic responses have been considered to be symmetrical between the Earth’s hemispheres. The other components of the photoperiod (dSR, dSS and dSSR) are mainly caused by the variable orbital angular speed of the ellipsoidal orbit around the sun. Consequently, they occur simultaneously at every latitude, shifting by six months between the cropping seasons in the two hemispheres. (Fig. 1B&C). Of the 77 early- to mid-early sorghum accessions tested in Madagascar, 43% exhibited significantly delayed heading times when sown in November–December. (Fig. 2A, Supp. Table 1). Similarly, 90% of the varieties flowered significantly later in Argentina than in the Northern Hemisphere (Fig. S4). This possible change was anticipated in response to the greater daily variations in sunrise and sunset times in December-January, when the Earth is at the perihelion of its orbit, i.e. at its closest distance to the Sun. This astronomical phase resulted in a prolonged PI in numerous sorghum varieties sown in the Northern Hemisphere from November to January (Bezot 1963; Miller *et al*. 1968; Clerget *et al*. 2021). A similar effect was therefore possible on sowings done at the onset of the rainy season, November to January, in the Southern Hemisphere. This effect has been clearly demonstrated in the two experiments presented in this paper. Only one of the three teams asked about such observations responded positively, which is understandable. Breeders carry out adaptation trials to test the adaptability of genetic materials to new environments. Consequently, non-adapted materials are discarded immediately without trace. It is fortunate that such a trial was conducted in Madagascar and in Argentina some years ago, and that the breeders still remember the unexpectedly long heading time of many cultivars.

On another hand, 17% of the cultivars were significantly earlier in Madagascar than in the Northern Hemisphere. All of these cultivars were early and all headed between 49 and 60 days after sowing in Madagascar, which is the shortest observed duration. Photoperiod-insensitive varieties such as BF201 typically exhibit this earlier heading time. This pattern has been observed for the first time. This suggests that, despite the long days, the photoperiod (i.e. dSR and dSS) at the end of the PIP was favourable for immediate panicle initiation in this group of cultivars.

Finally, the phenological model proposed in this paper suggests that the unusual delay in panicle initiation observed in four Tanzanian and one Ethiopian variety sown in November and December in Bamako may be due to their high sensitivity to dSR and dSS in their Southern Hemisphere origin.

### Several latent responses to photoperiod in each sorghum plant

The by-plant analysis of the unique dataset of durations to PI, recorded over six complete years for the CSM 335 variety, showed a significant year effect, particularly for sowings from November to February (Table 1, Fig. 3). Previously, the date of PI had been estimated on plot level, using data from all plants, and only the longer durations had been considered for sowings from November to February. This approach was influenced by previous papers on plant monthly sowings, which did not report similar observations (Miller *et al*. 1968) or did not consider them (Carberry *et al*. 2001). However, the presence of two to five groups of annual means of duration to PI in January-February and August-December sowings of CSM 335 that are significantly distinct could not be overlooked. They demonstrated that plants of this variety exhibit four potential, repeatable photoperiodic responses, with one being chosen in response to the environmental factors at the onset of the PSP. Series of 24 consecutive months of sowing were insufficient to reveal the four responses, and were instead interpreted based on the results obtained from CSM 335 (76 months), IRAT 174 (48 months) and Sariaso 10 (48 months).

CSM 335 is a streamlined Malian ecotype that was released in 2002 and has been maintained by breeding programmes through selfing ever since, so residual hidden inter-line variability could exist. However, with regard to flowering time, variability was strictly interannual for sowings from November to February, except in one plant out of 233, which statistically excludes inter-line variability. Conversely, the hypothesis that each sorghum line exhibits up to four potential photoperiodic responses affecting flowering time is supported by the presence of large numbers of duplicated genes in sorghum (Paterson *et al*. 2009). The first polyploidisation event occurred around 70 million years ago and affected almost all species in the Poaceae family. When sorghum and rice diverged 20 million years ago, only two copies of 30% of the duplicate genes had been retained. Due to their adaptive importance, two copies of genes in the flowering pathway may have been among this 30%. This was followed later by tandem duplications.

These four responses justified in hindsight why the Photoperiod Model 2.0 previously accounted for two seasonal responses: from June to September, when Kt1 is expressed; and from September to June, which includes two successive qualitative responses. (Clerget *et al*. 2021). Above all, they transformed the interpretation of the photoperiod response throughout the year. Rather than searching for a single response that could capture all the yearly variations, they facilitated a new and fruitful interpretation by combining alternative, independent responses to dSR/dSS rather than DL.

#### Internal coincidence model set at plant emergence in highly photoperiod sensitive sorghum

The identification of a linear relationship between dSR and dSS at the onset of the PSP and dSR+dSS at panicle initiation in the qualitative responses of late photoperiodic sorghum varieties confirmed the initial intuition. This leaded to hypothesize that the qualitative photoperiodic response resulted from an internal coincidence instead of a cumulative process as for the quantitative response. Indeed, according to Saunders (2021), four main models are currently used to explain the relationship between changes in daylength and the induction of events (e.g. diapause or flowering) in animals and plants. The second model is based on internal coincidence between two oscillators, one set at dawn and one at dusk, whose specific phasic relationship generates the inductive cue. While it fits our hypothesis, it needs to be complemented by an initial setting that has not previously been envisaged.

In insects, specific neurons in the brain of Drosophila melanogaster constitute morning and evening clocks (Menegazzi *et al*. 2020). In Arabidopsis thaliana and rice plants, the circadian clock gene network and morning and evening oscillators are now well characterised (Su *et al*. 2021; Michael 2022). This network comprises a central circadian oscillator involving two feedback-regulated genes whose transcription activity peaks in the morning and a series of eight genes whose activity peaks at specific times during the day and night. The activity of these ten genes is regulated by a series of photoreceptors that perceive light radiation between UV-B and far-red wavelengths, and which consequently detect dawn and dusk. The shoot apex is the focal point for synchronising all the cell oscillators (Takahashi *et al*. 2015). It is where plastochron is set at plant emergence and triggers new leaf initiation at regular time intervals (Brown *et al*. 2026). Thus, there is physiological support for an internal coincidence model involving two oscillators that generate the inductive cue when they reach exact phases set at the time of plant emergence.

### Solar photoperiod and daylength are not synonyms

Daylength is one of the components of the solar photoperiod, resulting from the time between sunrise and sunset. Plant cells use these last two components to set up the circadian clock (Su *et al*. 2021; Michael 2022). The yearly dynamics of sunrise and sunset times are much more complex (and consequently informative) than daylength. In a previous paper, we demonstrated that a model incorporating dSR, dSS and daylength could accurately predict the monthly duration to PI of sorghum crops over the course of a year at contrasting latitudes, a feat that no previous model had achieved (Clerget *et al*. 2021). In this paper, we hypothesized the coexistence of several photoperiodic responses in each plant, each acting during a different period of the year. Then, we demonstrated that the dSR and dSS factors alone are sufficient to accurately model these photoperiodic responses at a single latitude, provided that two sets of parameters are used: one above and one below 12 hours of daylight for the quantitative response. DL is also required to adjust the qualitative responses to variations in latitude. Both later model adjustments suggest that daylength plays a minor role in the flowering time of sorghum crops. Further evidence is needed to confirm these new hypotheses, either from field research or, more likely, from genetic analyses.

Thus, in the wild, plants would primarily depend on dSR and dSS rather than daylength for their seasonal photoperiodic reproduction. This results in similar responses from the equator to the polar circles. Conversely, in growth chambers, specific responses in flowering time to constant daylengths are observed and are related to the photoperiod sensitivity of the plants being tested. These responses demonstrate how plants adapt to an absence of natural synchronisation under a solar photoperiod. Similar dawn and dusk dynamics to those observed in the field can now be programmed in growth chambers, as described by (Goymann *et al*. 2012). This should result in observations and interpretations that are closer to what plants experience in the wild.

Lastly, the current conclusions concern the response of the duration to floral initiation to the natural photoperiod in the *Sorghum bicolor* species. As this species shares a similar genetic background to all other Poaceae species, and as the photoperiodic genetic pathways have remained largely unchanged throughout evolution, the hypotheses developed in sorghum should also be valid for the entire Poaceae family, and potentially beyond (Paterson *et al*. 2009). They should be applicable to both temperate and tropical species, and could be useful when flowering time is a crucial criterion for adaptation, as with temperate cereal crops. Indeed, sowing a mid-highly photoperiod-sensitive winter wheat variety between September and December would result in panicle initiation being triggered between 21 December (the winter solstice) and early February. This expected pattern mirrors the response of the CSM 335 short-day sorghum variety, when sown from March to June in the northern tropics (Fig. 6A). More photoperiod-sensitive wheat varieties when sown between September and December would initiate panicles later, in February or March.

## SUPPLEMENTARY DATA

Figure S1 Monthly mean temperatures season in five locations where the duration to flowering of sorghum crops is compared. Figure S2 Segmented linear regression of appeared leaf number on initiated leaf number. Figure S3 Monthly temperatures recorded in Samanko, Mali, from 2000 to 2009.

Figure S4 Duration to flowering of 27 cultivars observed in Argentina and in the northern hemisphere.

Table S1 List and durations to heading of 77 sorghum cultivars tested in Madagascar. Table S2 List of the 28 sorghum varieties monthly sown in Samanko, Mali. Table S3 Duration to flowering of 27 cultivars in Argentina and in the northern hemisphere. Table S4 ANOVAs of monthly data on the duration of panicle initiation in the 28 varieties. Table S5 Optimised parameters of the four photoperiodic responses without daylength effect for 28 varieties grown in Samanko. Table S6 Optimised parameters of the four photoperiodic responses with daylength effect in the quantitative responses for 28 varieties grown in Samanko. Table S7 Optimised parameters of the four photoperiodic responses for three varieties grown in Samanko and in Montpellier.

## DATA AVAILABILITY

**Clerget, B, Sidibe M.** 2026. Sorghum development and phenology data, Samanko, Mali, 2000-2009 [Data set]. Zenodo. https://doi.org/10.5281/zenodo.19354278

## AUTHOR CONTRIBUTIONS

BC, KvB, GT: conceptualisation, resources, supervision, editing; MS, VR, DO: experimental management, data management and curation; BC: data analysis, writing.

## AI ASSISTANCE

DeepSeek was used to generate the R scripts of the research model. DeepL-write was used to improve the English language.

## ACKNOWLEDGEMENTS

We would like to thank Denis Cornet from CIRAD for his help with R modelling using artificial intelligence, and Michael Dingkuhn, who has now retired from CIRAD, for his valuable comments on the draft paper. We are also grateful to Eva Weltzien and Fred Rattunde from ICRISAT for their invaluable support during the experimental period in Bamako.

