## Supplemental Table S4 for "Consequence and further analysis of the minor role of daylength in the floral initiation of sorghum crops"

### Up to four photoperiodic responses co-exist in sorghum plants and an internal phasic coincidence set at plant emergence trigger floral transition in absolute photoperiod-sensitive varieties

**Table S3:** Monthly data distribution with normality tests, and ANOVA of the year effect on the duration to panicle initiation in the 28 varieties.

##### Variety : Blanc Augaradeba

| **Statistic** | **Jan** | **Feb** | **Mar** | **Apr** | **May** | **Jun** | **Jul** | **Aug** | **Sep** | **Oct** | **Nov** | **Dec** |
|---|---|---|---|---|---|---|---|---|---|---|---|---|
| Mean | 50 | 183 | 170 | 150 | 111 | 91 | 70 | 56 | 42 | 38 | 46 | 35 |
| Std | 6 | 25 | 15 | 17 | 17 | 8 | 9 | 4 | 3 | 4 | 13 | 8 |
| N | 30 | 30 | 30 | 20 | 20 | 38 | 19 | 20 | 20 | 20 | 30 | 26 |
| DISTRIBUTION NORMALITY TESTS | | | | | | | | | | | | |
| Shapiro-Wilk | ns | ns | ns | *** | * | ns | ns | ns | ns | ns | ns | ** |
| Kolmogorov-Smirnov | ns | ns | ns | ** | ns | ns | ns | ns | ns | ns | ns | ns |
| Cramer-von Mises | ns | ns | ns | *** | * | ns | ns | ns | ns | ns | ns | ns |
| Anderson-Darling | ns | ns | ns | *** | * | ns | ns | ns | ns | ns | ns | * |
| ANOVA | | | | | | | | | | | | |
| Year Effect | *** | ns | ns | ns | ns | *** | ns | ns | * | ** | *** | *** |
| Yearly means | | | | | | | | | | | | |
| 2002 | | | | | | | | | | | 32 c | 43 a |
| 2003 | 45 b | 179 a | 179 a | 150 a | 112 a | 86 b | 68 a | 55 a | 40 b | 36 b | 60 a | 38 a |
| 2004 | 51 a | 184 a | 169 ab | 151 a | 110 a | 96 a | 73 a | 58 a | 43 a | 40 a | 46 b | 27 b |
| 2005 | 54 a | 185 a | 163 b | | | | | | | | | |

##### Variety : CGM 19

| **Statistic** | **Jan** | **Feb** | **Mar** | **Apr** | **May** | **Jun** | **Jul** | **Aug** | **Sep** | **Oct** | **Nov** | **Dec** |
|---|---|---|---|---|---|---|---|---|---|---|---|---|
| Mean | 57 | 83 | 99 | 87 | 67 | 51 | 53 | 36 | 30 | 33 | 70 | 56 |
| Std | 12 | 39 | 9 | 7 | 4 | 4 | 4 | 5 | 5 | 7 | 11 | 7 |
| N | 20 | 20 | 20 | 10 | 10 | 40 | 20 | 11 | 17 | 10 | 12 | 20 |
| DISTRIBUTION NORMALITY TESTS | | | | | | | | | | | | |
| Shapiro-Wilk | ** | *** | ns | * | ns | ** | *** | ns | * | * | ns | ns |
| Kolmogorov-Smirnov | * | *** | ns | ns | * | *** | ** | * | * | * | ns | ns |
| Cramer-von Mises | *** | *** | ns | * | ns | ** | ** | ns | ** | * | ns | ns |
| Anderson-Darling | *** | *** | ns | * | ns | ** | ** | ns | ** | * | ns | ns |
| ANOVA | | | | | | | | | | | | |
| Year Effect | *** | *** | * | | | *** | ns | ns | *** | *** | ns | ns |
| Yearly means | | | | | | | | | | | | |
| 2003 | | | | | | 48 b | 52 a | 32 a | 27 b | 31 b | 78 a | 55 a |
| 2004 | 68 a | 46 b | 94 b | 87 | 67 | 54 a | 53 a | 36 a | 34 a | 51 a | 68 a | 57 a |
| 2005 | 46 b | 120 a | 104 a | | | | | | | | | |

##### Variety : CS02 Mdg

| **Statistic** | **Jan** | **Feb** | **Mar** | **Apr** | **May** | **Jun** | **Jul** | **Aug** | **Sep** | **Oct** | **Nov** | **Dec** |
|---|---|---|---|---|---|---|---|---|---|---|---|---|
| Mean | 58 | 152 | 145 | 124 | 102 | 79 | 68 | 51 | 39 | 31 | 56 | 44 |
| Std | 26 | 13 | 13 | 10 | 4 | 6 | 8 | 6 | 4 | 5 | 16 | 10 |
| N | 30 | 30 | 30 | 20 | 20 | 40 | 19 | 20 | 20 | 20 | 30 | 30 |
| DISTRIBUTION NORMALITY TESTS | | | | | | | | | | | | |
| Shapiro-Wilk | *** | ns | ns | * | ns | ns | * | ns | *** | ns | ns | ns |
| Kolmogorov-Smirnov | *** | ns | ns | * | ns | ns | ns | ns | * | ns | ns | ns |
| Cramer-von Mises | *** | ns | ns | * | ns | ns | ** | ns | ** | ns | ns | ns |
| Anderson-Darling | *** | ns | ns | * | ns | ns | ** | ns | ** | ns | ns | ns |
| ANOVA | | | | | | | | | | | | |
| Year Effect | *** | * | *** | *** | ns | *** | *** | ns | ns | ns | *** | ** |
| Yearly means | | | | | | | | | | | | |
| 2002 | | | | | | | | | | | 46 b | 47 a |
| 2003 | 43 b | 147 b | 133 b | 132 a | 104 a | 74 b | 61 b | 52 a | 37 a | 30 a | 73 a | 48 a |
| 2004 | 89 a | 161 a | 148 a | 116 b | 101 a | 83 a | 76 a | 51 a | 40 a | 32 a | 48 b | 36 b |
| 2005 | 42 b | 147 b | 155 a | | | | | | | | | |

#### Variety : CSM 335

| **Statistic** | **Jan** | **Feb** | **Mar** | **Apr** | **May** | **Jun** | **Jul** | **Aug** | **Sep** | **Oct** | **Nov** | **Dec** |
|---|---|---|---|---|---|---|---|---|---|---|---|---|
| Mean | 65 | 99 | 107 | 93 | 76 | 60 | 50 | 38 | 28 | 28 | 64 | 62 |
| Std | 29 | 38 | 11 | 11 | 7 | 9 | 7 | 5 | 5 | 6 | 31 | 18 |
| N | 65 | 67 | 62 | 51 | 49 | 108 | 62 | 58 | 55 | 44 | 47 | 54 |
| DISTRIBUTION NORMALITY TESTS | | | | | | | | | | | | |
| Shapiro-Wilk | *** | *** | * | ns | ns | *** | ns | ns | ns | ** | * | ** |
| Kolmogorov-Smirnov | ns | *** | ns | ns | ns | *** | ns | ns | * | ** | ns | ns |
| Cramer-von Mises | * | *** | ns | ns | ns | *** | ns | ns | ns | ** | * | * |
| Anderson-Darling | ** | *** | ns | ns | ns | *** | ns | ns | ns | ** | * | * |
| ANOVA | | | | | | | | | | | | |
| Year Effect | *** | *** | *** | * | ** | *** | *** | *** | *** | *** | *** | *** |
| Yearly means | | | | | | | | | | | | |
| 2000 | | | | | | | 48 bc | 36 b | 22 c | 28 b | 84 b | 87 a |
| 2001 | 115 a | 130 a | 96 c | 96 ab | 71 b | 62 ab | 44 c | 44 a | 33 a | 35 a | 50 d | 70 b |
| 2002 | 71 b | 110 b | 109 ab | 99 a | 79 ab | 62 ab | 53 ab | 39 b | | | 26 e | 45 c |
| 2003 | 34 d | 58 c | 104 bc | 83 b | 74 b | 50 c | 51 abc | 38 b | 27 b | 25 b | 65 c | 48 c |
| 2004 | 39 cd | 40 d | 113 ab | 90 ab | 74 b | 60 b | 57 a | 34 b | 33 a | 27 b | | |
| 2005 | 49 c | 132 a | 111 ab | | | | | | | | | |
| 2006 | | | | | | | | | | | | 56 c |
| 2007 | 80 b | 130 a | 117 a | 101 a | 85 a | 54 c | 49 bc | 33 b | 27 b | 24 b | 111 a | 68 b |
| 2008 | 68 b | 102 b | 104 bc | 91 ab | 80 ab | 67 a | 46 bc | 36 b | 26 b | 36 a | | |

#### Variety : CSM 63-E

| **Statistic** | **Jan** | **Feb** | **Mar** | **Apr** | **May** | **Jun** | **Jul** | **Aug** | **Sep** | **Oct** | **Nov** | **Dec** |
|---|---|---|---|---|---|---|---|---|---|---|---|---|
| Mean | 66 | 45 | 42 | 41 | 33 | 29 | 33 | 28 | 26 | 22 | 83 | 60 |
| Std | 9 | 7 | 5 | 7 | 4 | 3 | 5 | 4 | 5 | 2 | 20 | 14 |
| N | 20 | 19 | 20 | 20 | 20 | 20 | 18 | 20 | 20 | 20 | 20 | 20 |
| DISTRIBUTION NORMALITY TESTS | | | | | | | | | | | | |
| Shapiro-Wilk | ns | ns | ns | ** | ns | ns | * | ns | ** | ns | ** | ns |
| Kolmogorov-Smirnov | ns | * | ns | * | ns | * | * | ns | ** | ns | ** | * |
| Cramer-von Mises | ns | * | ns | ** | ns | ns | ** | ns | ** | ns | ** | * |
| Anderson-Darling | ns | * | ns | ** | ns | ns | ** | ns | ** | ns | *** | * |
| ANOVA | | | | | | | | | | | | |
| Year Effect | ns | *** | *** | *** | *** | *** | ns | ns | *** | ns | *** | *** |
| Yearly means | | | | | | | | | | | | |
| 2002 | | | | | | | | | | | 64 b | 50 b |
| 2003 | 63 a | 52 a | 39 b | 48 a | 36 a | 26 b | 31 a | 28 a | 22 b | 21 a | 102 a | 70 a |
| 2004 | 68 a | 39 b | 45 a | 34 b | 30 b | 31 a | 35 a | 27 a | 30 a | 22 a | | |

#### Variety : Farafara 17

| **Statistic** | **Jan** | **Feb** | **Mar** | **Apr** | **May** | **Jun** | **Jul** | **Aug** | **Sep** | **Oct** | **Nov** | **Dec** |
|---|---|---|---|---|---|---|---|---|---|---|---|---|
| Mean | 47 | 160 | 133 | 126 | 96 | 80 | 66 | 47 | 32 | 27 | 54 | 43 |
| Std | 12 | 21 | 15 | 22 | 9 | 10 | 4 | 4 | 3 | 3 | 14 | 16 |
| N | 28 | 29 | 30 | 20 | 20 | 40 | 20 | 20 | 19 | 18 | 27 | 25 |
| DISTRIBUTION NORMALITY TESTS | | | | | | | | | | | | |
| Shapiro-Wilk | ** | ns | ns | ** | ** | *** | ns | * | ns | ns | ** | *** |
| Kolmogorov-Smirnov | ** | ns | ns | ** | *** | *** | ns | ** | ns | ns | * | ** |
| Cramer-von Mises | *** | ns | ns | *** | ** | *** | ns | ** | ns | ns | *** | *** |
| Anderson-Darling | *** | ns | ns | *** | ** | *** | ns | * | ns | ns | *** | *** |
| ANOVA | | | | | | | | | | | | |
| Year Effect | *** | ns | *** | * | ns | ns | * | * | * | ns | *** | *** |
| Yearly means | | | | | | | | | | | | |
| 2002 | | | | | | | | | | | 38 c | 47 b |
| 2003 | 39 b | 151 a | 120 c | 137 a | 92 a | 81 a | 64 b | 49 a | 30 b | 26 a | 67 a | 59 a |
| 2004 | 64 a | 165 a | 133 b | 115 b | 99 a | 80 a | 68 a | 45 b | 34 a | 28 a | 59 b | 25 c |
| 2005 | 42 b | 166 a | 147 a | | | | | | | | | |

#### Variety : IBS 19

| Statistic | Jan | Feb | Mar | Apr | May | Jun | Jul | Aug | Sep | Oct | Nov | Dec |
|---|---|---|---|---|---|---|---|---|---|---|---|---|
| Mean | 187 | 170 | 142 | 133 | 109 | 85 | 64 | 63 | 44 | 51 | 153 | 206 |
| Std | 26 | 23 | 22 | 12 | 17 | 14 | 5 | 5 | 4 | 21 | 72 | 30 |
| N | 20 | 18 | 16 | 15 | 18 | 18 | 20 | 15 | 16 | 15 | 11 | 17 |
| DISTRIBUTION NORMALITY TESTS | | | | | | | | | | | | |
| Shapiro-Wilk | ** | ** | * | * | *** | * | ns | ns | ns | ns | ns | * |
| Kolmogorov-Smirnov | ** | * | ns | ** | ** | ** | ns | ns | ns | * | ns | ** |
| Cramer-von Mises | ** | ** | ns | ** | *** | * | ns | ns | ns | * | ns | * |
| Anderson-Darling | ** | ** | * | ** | *** | * | ns | ns | ns | * | ns | * |
| ANOVA | | | | | | | | | | | | |
| Year Effect | ns | ns | ns | ns | * | ns | ns | ns | * | *** | *** | ns |
| Yearly means | | | | | | | | | | | | |
| 2006 | | | | | | | | | | | | 209 a |
| 2007 | 180 a | 173 a | 151 a | 131 a | 117 a | 80 a | 62 a | 65 a | 42 b | 36 b | 211 a | 205 a |
| 2008 | 195 a | 166 a | 136 a | 134 a | 101 b | 90 a | 65 a | 61 a | 47 a | 73 a | 83 b | |

#### Variety : IBS 30

| Statistic | Jan | Feb | Mar | Apr | May | Jun | Jul | Aug | Sep | Oct | Nov | Dec |
|---|---|---|---|---|---|---|---|---|---|---|---|---|
| Mean | 179 | 176 | 150 | 133 | 108 | 88 | 63 | 54 | 39 | 41 | 116 | 135 |
| Std | 38 | 17 | 16 | 13 | 12 | 4 | 3 | 16 | 3 | 7 | 55 | 70 |
| N | 18 | 40 | 16 | 18 | 13 | 40 | 20 | 14 | 18 | 13 | 14 | 16 |
| DISTRIBUTION NORMALITY TESTS | | | | | | | | | | | | |
| Shapiro-Wilk | ** | ** | ns | ** | *** | ns | ns | ** | ns | ns | * | * |
| Kolmogorov-Smirnov | ** | ns | * | *** | ** | ns | ns | *** | ** | ns | * | * |
| Cramer-von Mises | ** | * | * | ** | *** | ns | ns | *** | * | ns | * | * |
| Anderson-Darling | ** | * | * | *** | *** | ns | ns | *** | * | ns | * | ** |
| ANOVA | | | | | | | | | | | | |
| Year Effect | ns | * | ns | ns | ns | *** | ** | ns | ns | * | ** | *** |
| Yearly means | | | | | | | | | | | | |
| 2006 | | | | | | | | | | | | 197 a |
| 2007 | 176 a | 181 a | 152 a | 129 a | 110 a | 85 b | 61 b | 50 a | 39 a | 37 b | 156 a | 73 b |
| 2008 | 182 a | 170 b | 149 a | 136 a | 106 a | 90 a | 65 a | 57 a | 40 a | 46 a | 77 b | |

#### Variety : IBS 40

| Statistic | Jan | Feb | Mar | Apr | May | Jun | Jul | Aug | Sep | Oct | Nov | Dec |
|---|---|---|---|---|---|---|---|---|---|---|---|---|
| Mean | 187 | 174 | 153 | 133 | 103 | 76 | 58 | 54 | 37 | 82 | 129 | 180 |
| Std | 26 | 12 | 11 | 11 | 11 | 10 | 7 | 11 | 5 | 38 | 60 | 64 |
| N | 18 | 38 | 18 | 15 | 19 | 40 | 20 | 14 | 19 | 9 | 12 | 17 |
| DISTRIBUTION NORMALITY TESTS | | | | | | | | | | | | |
| Shapiro-Wilk | *** | ** | * | ns | ns | *** | *** | ns | ns | ns | ** | * |
| Kolmogorov-Smirnov | * | *** | ns | ns | ns | *** | ** | ns | ns | ns | *** | * |
| Cramer-von Mises | *** | *** | * | ns | ns | *** | *** | ns | ns | ns | *** | * |
| Anderson-Darling | *** | *** | * | ns | ns | *** | *** | ns | ns | ns | *** | * |
| ANOVA | | | | | | | | | | | | |
| Year Effect | ns | ns | ns | ns | ** | * | ns | ns | * | ** | ** | ns |
| Yearly means | | | | | | | | | | | | |
| 2006 | | | | | | | | | | | | 170 a |
| 2007 | 188 a | 176 a | 157 a | 127 a | 109 a | 80 a | 58 a | 58 a | 35 b | 37 b | 194 a | 192 a |
| 2008 | 186 a | 172 a | 149 a | 137 a | 96 b | 73 b | 58 a | 51 a | 40 a | 104 a | 97 b | |

#### Variety : IBS 582

| **Statistic** | **Jan** | **Feb** | **Mar** | **Apr** | **May** | **Jun** | **Jul** | **Aug** | **Sep** | **Oct** | **Nov** | **Dec** |
|---|---|---|---|---|---|---|---|---|---|---|---|---|
| Mean | 187 | 181 | 152 | 134 | 113 | 83 | 61 | 55 | 39 | 71 | 149 | 164 |
| Std | 18 | 9 | 17 | 10 | 9 | 9 | 5 | 6 | 5 | 43 | 56 | 61 |
| N | 17 | 36 | 19 | 19 | 12 | 36 | 19 | 18 | 18 | 11 | 13 | 14 |
| DISTRIBUTION NORMALITY TESTS | | | | | | | | | | | | |
| Shapiro-Wilk | ns | *** | ** | *** | ns | ns | ns | ns | ns | * | ns | ** |
| Kolmogorov-Smirnov | ns | *** | * | * | ns | * | ns | ns | ns | * | * | ** |
| Cramer-von Mises | ns | *** | *** | *** | ns | ns | ns | ns | ns | * | ns | ** |
| Anderson-Darling | ns | *** | *** | *** | ns | ns | ns | ns | ns | * | ns | ** |
| ANOVA | | | | | | | | | | | | |
| Year Effect | ns | ns | ns | ns | ns | * | ** | ns | ns | *** | ** | ns |
| Yearly means | | | | | | | | | | | | |
| 2006 | | | | | | | | | | | | 142 a |
| 2007 | 182 a | 182 a | 160 a | 131 a | 114 a | 80 b | 58 b | 56 a | 40 a | 35 b | 180 a | 186 a |
| 2008 | 193 a | 179 a | 146 a | 137 a | 113 a | 87 a | 64 a | 54 a | 38 a | 114 a | 99 b | |

#### Variety : IRAT 174

| **Statistic** | **Jan** | **Feb** | **Mar** | **Apr** | **May** | **Jun** | **Jul** | **Aug** | **Sep** | **Oct** | **Nov** | **Dec** |
|---|---|---|---|---|---|---|---|---|---|---|---|---|
| Mean | 47 | 33 | 31 | 87 | 77 | 63 | 50 | 35 | 22 | 22 | 67 | 64 |
| Std | 10 | 3 | 6 | 4 | 7 | 5 | 5 | 5 | 3 | 4 | 22 | 13 |
| N | 40 | 40 | 38 | 39 | 36 | 80 | 45 | 46 | 37 | 30 | 36 | 34 |
| DISTRIBUTION NORMALITY TESTS | | | | | | | | | | | | |
| Shapiro-Wilk | ** | ns | *** | ns | ns | ** | *** | ns | ** | ** | *** | ns |
| Kolmogorov-Smirnov | * | ns | *** | ns | ns | ** | *** | ns | ** | ** | *** | ns |
| Cramer-von Mises | ** | ns | *** | ns | ns | * | *** | ns | ** | ** | *** | ns |
| Anderson-Darling | ** | ns | *** | ns | ns | ** | *** | ns | ** | ** | *** | ns |
| ANOVA | | | | | | | | | | | | |
| Year Effect | *** | *** | *** | ns | *** | *** | ns | *** | *** | *** | *** | *** |
| Yearly means | | | | | | | | | | | | |
| 2000 | | | | | | | 53 a | 34 bc | 22 b | 19 b | 72 b | 79 a |
| 2001 | 59 a | 34 b | 28 b | 86 a | 72 b | 62 b | 52 a | 36 b | 27 a | 22 b | 50 c | 62 b |
| 2002 | 42 c | 29 d | 27 b | 87 a | 82 a | 66 a | 51 a | 42 a | | | | |
| 2006 | | | | | | | | | | | | 56 b |
| 2007 | 51 b | 36 a | 40 a | 89 a | 72 b | 62 b | 50 a | 33 bc | 20 b | 19 b | 92 a | 54 b |
| 2008 | 34 d | 32 c | 28 b | 88 a | 82 a | 61 b | 48 a | 31 c | 21 b | 27 a | 50 c | |

#### Variety : IRAT 204

| **Statistic** | **Jan** | **Feb** | **Mar** | **Apr** | **May** | **Jun** | **Jul** | **Aug** | **Sep** | **Oct** | **Nov** | **Dec** |
|---|---|---|---|---|---|---|---|---|---|---|---|---|
| Mean | 21 | 22 | 19 | 25 | 29 | 29 | 43 | 38 | 27 | 23 | 20 | 24 |
| Std | 4 | 6 | 2 | 5 | 1 | 2 | 11 | 6 | 3 | 2 | 2 | 7 |
| N | 30 | 30 | 29 | 18 | 20 | 20 | 17 | 17 | 20 | 17 | 30 | 20 |
| DISTRIBUTION NORMALITY TESTS | | | | | | | | | | | | |
| Shapiro-Wilk | ** | ** | ns | ns | ns | ns | * | ns | ns | ns | ** | ** |
| Kolmogorov-Smirnov | ** | *** | ns | ns | ns | ns | * | ns | ns | ns | ** | * |
| Cramer-von Mises | *** | *** | ns | ns | ns | ns | * | ns | ns | ns | ** | * |
| Anderson-Darling | *** | *** | ns | ns | ns | ns | * | ns | ns | ns | ** | ** |
| ANOVA | | | | | | | | | | | | |
| Year Effect | *** | *** | ns | *** | ns | ns | ** | ns | *** | ** | ** | *** |
| Yearly means | | | | | | | | | | | | |
| 2002 | | | | | | | | | | | 18 b | 31 a |
| 2003 | 23 a | 18 b | 18 a | 30 a | 29 a | 29 a | 34 b | 37 a | 25 b | 22 b | 21 a | |
| 2004 | 17 b | 18 b | 19 a | 21 b | 30 a | 28 a | 50 a | 40 a | 29 a | 25 a | 20 a | 17 b |
| 2005 | 23 a | 29 a | 19 a | | | | | | | | | |

#### Variety : IS 11026

| **Statistic** | **Jan** | **Feb** | **Mar** | **Apr** | **May** | **Jun** | **Jul** | **Aug** | **Sep** | **Oct** | **Nov** | **Dec** |
|---|---|---|---|---|---|---|---|---|---|---|---|---|
| Mean | 129 | 121 | 95 | 79 | 77 | 59 | 59 | 55 | 47 | 40 | 107 | 97 |
| Std | 9 | 7 | 3 | 3 | 7 | 5 | 13 | 18 | 6 | 2 | 41 | 12 |
| N | 9 | 10 | 9 | 8 | 7 | 34 | 18 | 6 | 10 | 4 | 7 | 10 |
| DISTRIBUTION NORMALITY TESTS | | | | | | | | | | | | |
| Shapiro-Wilk | ns | ns | ns | ns | ns | ns | ns | ns | *** | ns | * | ns |
| Kolmogorov-Smirnov | ns | ns | ns | ns | ns | * | ns | ns | ** | | * | ns |
| Cramer-von Mises | ns | ns | ns | ns | | ns | ns | | *** | | | ns |
| Anderson-Darling | ns | ns | ns | ns | | ns | ns | | *** | | | ns |
| ANOVA | | | | | | | | | | | | |
| Year Effect | | | | | | ns | ** | | | | | |
| Yearly means | | | | | | | | | | | | |
| 2007 | | | | | | 59 a | 51 b | 55 | 47 | 40 | 107 | 97 |
| 2008 | 129 | 121 | 95 | 79 | 77 | 60 a | 67 a | | | | | |

#### Variety : IS 12169

| **Statistic** | **Jan** | **Feb** | **Mar** | **Apr** | **May** | **Jun** | **Jul** | **Aug** | **Sep** | **Oct** | **Nov** | **Dec** |
|---|---|---|---|---|---|---|---|---|---|---|---|---|
| Mean | 153 | 147 | 146 | 90 | 88 | 70 | 62 | 41 | 33 | 27 | 134 | 70 |
| Std | 37 | 18 | 5 | 2 | 7 | 6 | 11 | 8 | 6 | 2 | 49 | 4 |
| N | 10 | 17 | 10 | 9 | 9 | 26 | 18 | 5 | 10 | 5 | 10 | 9 |
| DISTRIBUTION NORMALITY TESTS | | | | | | | | | | | | |
| Shapiro-Wilk | ** | *** | *** | ns | ns | ns | ** | ns | * | ns | *** | ns |
| Kolmogorov-Smirnov | ** | ** | ** | ns | ns | ns | * | * | ** | ns | *** | ns |
| Cramer-von Mises | ** | *** | ** | ns | ns | ns | * | | * | | *** | ns |
| Anderson-Darling | ** | *** | *** | ns | ns | ns | ** | | * | | *** | ns |
| ANOVA | | | | | | | | | | | | |
| Year Effect | | | | | | * | ns | | | | | |
| Yearly means | | | | | | | | | | | | |
| 2007 | | | | | | 68 b | 59 a | 41 | 33 | 27 | 134 | 70 |
| 2008 | 153 | 147 | 146 | 90 | 88 | 72 a | 65 a | | | | | |

#### Variety : IS 15401

| **Statistic** | **Jan** | **Feb** | **Mar** | **Apr** | **May** | **Jun** | **Jul** | **Aug** | **Sep** | **Oct** | **Nov** | **Dec** |
|---|---|---|---|---|---|---|---|---|---|---|---|---|
| Mean | 33 | 187 | 170 | 134 | 96 | 95 | 73 | 52 | 34 | 26 | 29 | 37 |
| Std | 2 | 27 | 8 | 10 | 23 | 11 | 15 | 4 | 6 | 3 | 2 | 3 |
| N | 20 | 20 | 20 | 20 | 20 | 40 | 20 | 20 | 20 | 20 | 20 | 20 |
| DISTRIBUTION NORMALITY TESTS | | | | | | | | | | | | |
| Shapiro-Wilk | ns | ns | ns | * | ** | *** | ** | ns | * | ns | ns | ns |
| Kolmogorov-Smirnov | ns | ns | * | ** | *** | *** | * | ns | ** | ns | ns | ns |
| Cramer-von Mises | ns | ns | ns | * | *** | *** | ** | ns | * | ns | ns | * |
| Anderson-Darling | ns | ns | ns | * | *** | *** | ** | ns | * | ns | ns | ns |
| ANOVA | | | | | | | | | | | | |
| Year Effect | ns | ** | ns | ns | *** | *** | * | ns | *** | *** | ns | ** |
| Yearly means | | | | | | | | | | | | |
| 2002 | | | | | | | | | | | 28 a | 38 a |
| 2003 | 33 a | 203 a | 170 a | 131 a | 75 b | 86 b | 66 b | 54 a | 28 b | 23 b | 29 a | 35 b |
| 2004 | 33 a | 171 b | 170 a | 136 a | 118 a | 103 a | 80 a | 50 a | 40 a | 28 a | | |

#### Variety : IS 6351

| **Statistic** | **Jan** | **Feb** | **Mar** | **Apr** | **May** | **Jun** | **Jul** | **Aug** | **Sep** | **Oct** | **Nov** | **Dec** |
|---|---|---|---|---|---|---|---|---|---|---|---|---|
| Mean | 64 | 84 | 90 | 82 | 78 | 64 | 56 | 42 | 37 | 31 | 103 | 75 |
| Std | 10 | 4 | 7 | 6 | 5 | 5 | 9 | 7 | 1 | 4 | 9 | 6 |
| N | 10 | 10 | 6 | 10 | 9 | 20 | 18 | 6 | 10 | 3 | 10 | 10 |
| DISTRIBUTION NORMALITY TESTS | | | | | | | | | | | | |
| Shapiro-Wilk | * | ns | ns | ns | ns | ns | ns | ns | ns | ns | ns | ns |
| Kolmogorov-Smirnov | ns | ns | ns | ns | ns | ns | * | ns | *** | | ns | ns |
| Cramer-von Mises | * | ns | | ns | ns | ns | ns | | ** | | ns | ns |
| Anderson-Darling | * | ns | | ns | ns | ns | ns | | ** | | ns | ns |
| ANOVA | | | | | | | | | | | | |
| Year Effect | | | | | | ns | * | | | | | |
| Yearly means | | | | | | | | | | | | |
| 2007 | | | | | | 63 a | 51 b | 42 | 37 | 31 | 103 | 75 |
| 2008 | 64 | 84 | 90 | 82 | 78 | 64 a | 60 a | | | | | |

#### Variety : Kaura D-12

| **Statistic** | **Jan** | **Feb** | **Mar** | **Apr** | **May** | **Jun** | **Jul** | **Aug** | **Sep** | **Oct** | **Nov** | **Dec** |
|---|---|---|---|---|---|---|---|---|---|---|---|---|
| Mean | 40 | 102 | 139 | 115 | 102 | 81 | 60 | 42 | 27 | 22 | 39 | 38 |
| Std | 6 | 36 | 18 | 19 | 13 | 10 | 11 | 7 | 4 | 2 | 11 | 11 |
| N | 26 | 30 | 30 | 20 | 20 | 38 | 20 | 18 | 20 | 20 | 30 | 28 |
| DISTRIBUTION NORMALITY TESTS | | | | | | | | | | | | |
| Shapiro-Wilk | ns | ns | ns | ns | ns | ns | ** | * | * | ns | *** | ns |
| Kolmogorov-Smirnov | ns | ns | ns | ns | ns | ns | * | * | * | ** | * | ns |
| Cramer-von Mises | ns | ns | * | ns | ns | ns | ** | * | * | ** | *** | ns |
| Anderson-Darling | ns | ns | * | ns | ns | ns | ** | * | * | * | *** | ns |
| ANOVA | | | | | | | | | | | | |
| Year Effect | *** | * | ns | ns | ns | * | ns | * | *** | * | ns | *** |
| Yearly means | | | | | | | | | | | | |
| 2002 | | | | | | | | | | | 36 a | 42 b |
| 2003 | 35 b | 116 a | 144 a | 113 a | 98 a | 77 b | 59 a | 46 a | 24 b | 22 b | 43 a | 48 a |
| 2004 | 42 a | 79 b | 143 a | 117 a | 107 a | 84 a | 61 a | 39 b | 30 a | 23 a | 39 a | 26 c |
| 2005 | 44 a | 112 a | 131 a | | | | | | | | | |

#### Variety : Kende Bla

| **Statistic** | **Jan** | **Feb** | **Mar** | **Apr** | **May** | **Jun** | **Jul** | **Aug** | **Sep** | **Oct** | **Nov** | **Dec** |
|---|---|---|---|---|---|---|---|---|---|---|---|---|
| Mean | 37 | 64 | 63 | 55 | 52 | 41 | 35 | 28 | 21 | 30 | 34 | 26 |
| Std | 6 | 26 | 15 | 2 | 3 | 5 | 4 | 5 | 3 | 2 | 4 | 7 |
| N | 18 | 20 | 20 | 20 | 19 | 20 | 17 | 14 | 9 | 6 | 20 | 20 |
| DISTRIBUTION NORMALITY TESTS | | | | | | | | | | | | |
| Shapiro-Wilk | ns | ** | ** | ns | ns | * | ns | ns | ** | ns | * | * |
| Kolmogorov-Smirnov | * | ** | ** | ns | ns | * | ns | ns | ** | ns | * | ** |
| Cramer-von Mises | ns | *** | ** | ns | ns | ** | ns | ns | ** | | * | * |
| Anderson-Darling | ns | *** | ** | ns | ns | ** | ns | ns | ** | | * | * |
| ANOVA | | | | | | | | | | | | |
| Year Effect | *** | *** | *** | ** | ns | ** | ns | ns | ** | | ns | *** |
| Yearly means | | | | | | | | | | | | |
| 2003 | | | | 56 a | 52 a | 38 b | 33 a | 26 a | 20 b | 30 | 34 a | 32 a |
| 2004 | 41 a | 39 b | 50 b | 54 b | 52 a | 44 a | 36 a | 28 a | 28 a | | 34 a | 20 b |
| 2005 | 32 b | 88 a | 77 a | | | | | | | | | |

#### Variety : Macia

| **Statistic** | **Jan** | **Feb** | **Mar** | **Apr** | **May** | **Jun** | **Jul** | **Aug** | **Sep** | **Oct** | **Nov** | **Dec** |
|---|---|---|---|---|---|---|---|---|---|---|---|---|
| Mean | 28 | 37 | 31 | 35 | 36 | 36 | 40 | 48 | 33 | 29 | 38 | 44 |
| Std | 3 | 4 | 5 | 6 | 3 | 3 | 3 | 5 | 3 | 3 | 8 | 5 |
| N | 20 | 19 | 20 | 19 | 20 | 19 | 20 | 16 | 17 | 13 | 14 | 17 |
| DISTRIBUTION NORMALITY TESTS | | | | | | | | | | | | |
| Shapiro-Wilk | ns | ns | * | * | * | ns | ** | ns | * | ns | * | ns |
| Kolmogorov-Smirnov | ns | ns | ns | * | ns | ns | ** | ns | * | ns | ns | ns |
| Cramer-von Mises | ns | ns | ns | * | ns | ns | ** | ns | * | ns | * | ns |
| Anderson-Darling | ns | ns | ns | * | ns | ns | ** | ns | * | ns | * | ns |
| ANOVA | | | | | | | | | | | | |
| Year Effect | ns | ns | *** | *** | ns | ** | *** | ns | *** | ns | ns | * |
| Yearly means | | | | | | | | | | | | |
| 2006 | | | | | | | | | | | | 42 b |
| 2007 | 28 a | 38 a | 27 b | 30 b | 38 a | 34 b | 42 a | 48 a | 30 b | 30 a | 41 a | 47 a |
| 2008 | 29 a | 37 a | 35 a | 40 a | 35 a | 38 a | 38 b | 49 a | 35 a | 29 a | 34 a | |

#### Variety : Mtama

| Statistic | Jan | Feb | Mar | Apr | May | Jun | Jul | Aug | Sep | Oct | Nov | Dec |
|---|---|---|---|---|---|---|---|---|---|---|---|---|
| Mean | 43 | 45 | 50 | 52 | 50 | 46 | 44 | 41 | 26 | 26 | 49 | 49 |
| Std | 7 | 3 | 6 | 6 | 2 | 5 | 8 | 5 | 2 | 4 | 16 | 4 |
| N | 19 | 20 | 19 | 19 | 19 | 20 | 20 | 14 | 16 | 19 | 16 | 18 |
| DISTRIBUTION NORMALITY TESTS | | | | | | | | | | | | |
| Shapiro-Wilk | ns | ns | ns | ns | ns | * | *** | ns | * | ** | ** | ns |
| Kolmogorov-Smirnov | ns | ns | ns | ns | ns | ns | *** | ns | ns | * | ** | ns |
| Cramer-von Mises | ns | ns | ns | ns | ns | ns | *** | ns | ns | * | ** | ns |
| Anderson-Darling | ns | ns | ns | ns | ns | * | *** | ns | * | ** | ** | ns |
| ANOVA | | | | | | | | | | | | |
| Year Effect | ns | *** | *** | *** | ns | *** | ns | ns | ns | ** | *** | ns |
| Yearly means | | | | | | | | | | | | |
| 2006 | | | | | | | | | | | | 50 a |
| 2007 | 42 a | 48 a | 46 b | 47 b | 50 a | 42 b | 43 a | 42 a | 27 a | 24 b | 62 a | 48 a |
| 2008 | 45 a | 43 b | 55 a | 56 a | 49 a | 51 a | 46 a | 40 a | 25 a | 28 a | 33 b | |

#### Variety : Oueni

| Statistic | Jan | Feb | Mar | Apr | May | Jun | Jul | Aug | Sep | Oct | Nov | Dec |
|---|---|---|---|---|---|---|---|---|---|---|---|---|
| Mean | 51 | 62 | 95 | 76 | 63 | 52 | 45 | 30 | 29 | 30 | 72 | 50 |
| Std | 12 | 18 | 19 | 11 | 10 | 5 | 7 | 5 | 8 | 10 | 14 | 10 |
| N | 17 | 20 | 20 | 20 | 20 | 40 | 14 | 14 | 17 | 20 | 20 | 40 |
| DISTRIBUTION NORMALITY TESTS | | | | | | | | | | | | |
| Shapiro-Wilk | ** | ns | ns | ns | ns | *** | ns | ns | ns | *** | ns | ** |
| Kolmogorov-Smirnov | * | ns | ns | ns | ns | *** | ns | ns | ns | * | ns | ** |
| Cramer-von Mises | ** | ns | ns | ns | ns | *** | ns | ns | ns | ** | ns | ** |
| Anderson-Darling | ** | ns | ns | ns | ns | *** | ns | ns | ns | ** | ns | ** |
| ANOVA | | | | | | | | | | | | |
| Year Effect | ** | ns | * | ns | ns | ns | * | * | *** | ** | ** | *** |
| Yearly means | | | | | | | | | | | | |
| 2003 | | | | 73 a | 63 a | 51 a | 38 b | 34 a | 24 b | 24 b | 64 b | 57 a |
| 2004 | 59 a | 54 a | 86 b | 78 a | 63 a | 52 a | 48 a | 28 b | 37 a | 37 a | 80 a | 43 b |
| 2005 | 42 b | 69 a | 105 a | | | | | | | | | |

#### Variety : Sariaso 10

| Statistic | Jan | Feb | Mar | Apr | May | Jun | Jul | Aug | Sep | Oct | Nov | Dec |
|---|---|---|---|---|---|---|---|---|---|---|---|---|
| Mean | 48 | 49 | 52 | 60 | 57 | 50 | 44 | 41 | 28 | 26 | 47 | 51 |
| Std | 10 | 9 | 9 | 8 | 6 | 7 | 4 | 6 | 3 | 5 | 18 | 10 |
| N | 33 | 38 | 37 | 39 | 36 | 35 | 45 | 46 | 37 | 36 | 34 | 38 |
| DISTRIBUTION NORMALITY TESTS | | | | | | | | | | | | |
| Shapiro-Wilk | ns | ns | ns | *** | ns | ns | ns | ** | ns | ns | *** | ns |
| Kolmogorov-Smirnov | ns | ns | ns | ns | ns | ns | ns | ** | ns | ns | *** | ns |
| Cramer-von Mises | ns | ns | ns | * | * | ns | ns | *** | ns | ns | *** | * |
| Anderson-Darling | ns | ns | ns | * | * | ns | ns | *** | ns | ns | *** | * |
| ANOVA | | | | | | | | | | | | |
| Year Effect | *** | *** | *** | ns | * | *** | *** | *** | *** | *** | *** | ** |
| Yearly means | | | | | | | | | | | | |
| 2000 | | | | | | | 40 c | 44 a | 25 c | 24 b | 32 c | 54 ab |
| 2001 | 57 a | 52 ab | 57 a | 56 a | 57 ab | 49 b | 46 ab | 45 a | 32 a | 30 a | 42 b | 45 b |
| 2002 | 41 b | 40 c | 40 b | 57 a | 58 ab | 58 a | 47 a | 43 a | | | | |
| 2006 | | | | | | | | | | | | 58 a |
| 2007 | 56 a | 57 a | 52 a | 64 a | 60 a | 44 b | 44 ab | 31 c | 28 b | 22 b | 80 a | 47 b |
| 2008 | 41 b | 47 b | 57 a | 62 a | 52 b | 48 b | 43 bc | 39 b | 28 b | 29 a | 43 b | |

#### Variety : Short Kaura

| Statistic | Jan | Feb | Mar | Apr | May | Jun | Jul | Aug | Sep | Oct | Nov | Dec |
|---|---|---|---|---|---|---|---|---|---|---|---|---|
| Mean | 84 | 157 | 161 | 142 | 125 | 102 | 84 | 50 | 50 | 43 | 50 | 51 |
| Std | 21 | 27 | 16 | 12 | 16 | 8 | 10 | 14 | 2 | 4 | 4 | 14 |
| N | 28 | 30 | 30 | 20 | 20 | 40 | 19 | 10 | 10 | 10 | 10 | 30 |
| DISTRIBUTION NORMALITY TESTS | | | | | | | | | | | | |
| Shapiro-Wilk | ** | ns | * | ns | ** | ** | * | ns | ns | ns | ns | ** |
| Kolmogorov-Smirnov | ns | ns | ** | ns | * | *** | ** | ns | ns | ns | ns | ** |
| Cramer-von Mises | * | ns | ** | ns | ** | ** | * | ns | ns | * | ns | ** |
| Anderson-Darling | ** | ns | * | ns | ** | ** | * | ns | ns | * | ns | ** |
| ANOVA | | | | | | | | | | | | |
| Year Effect | *** | ns | ns | ns | ** | *** | ns | | | | | *** |
| Yearly means | | | | | | | | | | | | |
| 2002 | | | | | | | | | | | | 64 a |
| 2003 | 81 b | 156 a | 157 a | 140 a | 115 b | 94 b | 82 a | | | | | 56 b |
| 2004 | 107 a | 165 a | 163 a | 143 a | 136 a | 109 a | 84 a | 50 | 50 | 43 | 50 | 33 c |
| 2005 | 66 c | 152 a | 163 a | | | | | | | | | |

#### Variety : Sima

| Statistic | Jan | Feb | Mar | Apr | May | Jun | Jul | Aug | Sep | Oct | Nov | Dec |
|---|---|---|---|---|---|---|---|---|---|---|---|---|
| Mean | 37 | 46 | 52 | 58 | 54 | 46 | 46 | 40 | 26 | 26 | 45 | 52 |
| Std | 5 | 5 | 8 | 6 | 2 | 5 | 4 | 6 | 4 | 5 | 9 | 4 |
| N | 19 | 20 | 15 | 18 | 20 | 20 | 20 | 17 | 17 | 17 | 14 | 16 |
| DISTRIBUTION NORMALITY TESTS | | | | | | | | | | | | |
| Shapiro-Wilk | ns | ns | ns | ns | ns | * | * | * | ns | ** | ns | ns |
| Kolmogorov-Smirnov | ns | ns | ns | ns | ns | ns | ns | ns | ns | ns | ns | ns |
| Cramer-von Mises | ns | ns | ns | ns | ns | * | ns | * | ns | * | ns | ns |
| Anderson-Darling | ns | ns | ns | ns | ns | * | ns | * | ns | ** | ns | ns |
| ANOVA | | | | | | | | | | | | |
| Year Effect | ** | ** | *** | ** | ns | ns | ns | ns | ns | ** | ns | ns |
| Yearly means | | | | | | | | | | | | |
| 2006 | | | | | | | | | | | | 52 a |
| 2007 | 40 a | 49 a | 44 b | 63 a | 54 a | 44 a | 45 a | 40 a | 27 a | 23 b | 48 a | 51 a |
| 2008 | 34 b | 43 b | 58 a | 55 b | 53 a | 48 a | 46 a | 40 a | 24 a | 29 a | 39 a | |

#### Variety : Souroukoukou

| Statistic | Jan | Feb | Mar | Apr | May | Jun | Jul | Aug | Sep | Oct | Nov | Dec |
|---|---|---|---|---|---|---|---|---|---|---|---|---|
| Mean | 45 | 206 | 163 | 146 | 126 | 103 | 78 | 67 | 40 | 33 | 47 | 58 |
| Std | 7 | 20 | 14 | 13 | 14 | 5 | 17 | 10 | 6 | 1 | 11 | 13 |
| N | 17 | 20 | 20 | 20 | 16 | 40 | 20 | 20 | 10 | 10 | 20 | 20 |
| DISTRIBUTION NORMALITY TESTS | | | | | | | | | | | | |
| Shapiro-Wilk | ns | ns | ns | ** | ** | * | ns | ns | *** | ns | ns | ns |
| Kolmogorov-Smirnov | * | ns | ns | ns | * | ns | ns | ns | ** | ns | ns | * |
| Cramer-von Mises | ns | ns | ns | ns | * | ns | ns | ns | *** | ns | ns | * |
| Anderson-Darling | ns | ns | ns | * | * | ns | ns | ns | *** | ns | ns | * |
| ANOVA | | | | | | | | | | | | |
| Year Effect | ** | ns | ns | ns | ns | *** | ** | ns | | | * | *** |
| Yearly means | | | | | | | | | | | | |
| 2002 | | | | | | | | | | | 42 b | 69 a |
| 2003 | 42 b | 212 a | 162 a | 144 a | 125 a | 100 b | 89 a | 68 a | | | 51 a | 46 b |
| 2004 | 50 a | 200 a | 163 a | 148 a | 126 a | 106 a | 68 b | 66 a | 40 | 33 | | |

#### Variety : SSM12

| Statistic | Jan | Feb | Mar | Apr | May | Jun | Jul | Aug | Sep | Oct | Nov | Dec |
|---|---|---|---|---|---|---|---|---|---|---|---|---|
| Mean | 34 | 146 | 143 | 124 | 113 | 78 | 65 | 48 | 48 | 45 | 73 | 48 |
| Std | 3 | 19 | 24 | 23 | 19 | 14 | 17 | 15 | 9 | 9 | 12 | 4 |
| N | 19 | 11 | 16 | 19 | 16 | 36 | 18 | 9 | 14 | 13 | 15 | 16 |
| DISTRIBUTION NORMALITY TESTS | | | | | | | | | | | | |
| Shapiro-Wilk | ns | * | * | *** | *** | * | ns | ns | *** | * | ** | * |
| Kolmogorov-Smirnov | ns | * | ** | *** | *** | ns | ns | ns | *** | ns | * | * |
| Cramer-von Mises | ns | * | * | *** | *** | ns | ns | ns | ** | ns | ** | * |
| Anderson-Darling | ns | * | * | *** | *** | ns | ns | ns | *** | * | ** | * |
| ANOVA | | | | | | | | | | | | |
| Year Effect | ns | ns | ** | ** | ns | ** | * | ns | ns | *** | *** | *** |
| Yearly means | | | | | | | | | | | | |
| 2006 | | | | | | | | | | 48 a | 60 b | 44 b |
| 2007 | 34 a | 124 a | 156 a | 138 a | 113 a | 84 a | 55 b | 46 a | 45 a | 37 b | 81 a | 52 a |
| 2008 | 34 a | 148 a | 126 b | 108 b | 112 a | 72 b | 75 a | 65 a | 53 a | 54 a | | |

#### Variety : SSM1592

| Statistic | Jan | Feb | Mar | Apr | May | Jun | Jul | Aug | Sep | Oct | Nov | Dec |
|---|---|---|---|---|---|---|---|---|---|---|---|---|
| Mean | 83 | 143 | 166 | 131 | 110 | 87 | 63 | 58 | 48 | 49 | 86 | 56 |
| Std | 24 | 44 | 15 | 22 | 14 | 13 | 11 | 11 | 6 | 12 | 15 | 5 |
| N | 18 | 20 | 19 | 20 | 16 | 32 | 14 | 9 | 11 | 11 | 10 | 15 |
| DISTRIBUTION NORMALITY TESTS | | | | | | | | | | | | |
| Shapiro-Wilk | ns | ns | ** | *** | * | *** | ns | ns | ns | ns | * | ns |
| Kolmogorov-Smirnov | ns | ns | * | *** | ns | *** | ns | ns | ns | ns | ns | ** |
| Cramer-von Mises | ns | ns | * | *** | * | *** | ns | ns | ns | ns | ns | ns |
| Anderson-Darling | ns | ns | * | *** | * | *** | ns | ns | ns | ns | * | ns |
| ANOVA | | | | | | | | | | | | |
| Year Effect | ** | ns | * | ns | ns | ns | ns | ns | * | ** | | ns |
| Yearly means | | | | | | | | | | | | |
| 2006 | | | | | | | | | | 61 a | | 55 a |
| 2007 | 99 a | 144 a | 173 a | 138 a | 113 a | 88 a | 65 a | 58 a | 45 b | 36 b | 86 | 56 a |
| 2008 | 66 b | 142 a | 158 b | 125 a | 106 a | 86 a | 60 a | 60 a | 55 a | 54 a | | |

#### Variety : SSM29

| Statistic | Jan | Feb | Mar | Apr | May | Jun | Jul | Aug | Sep | Oct | Nov | Dec |
|---|---|---|---|---|---|---|---|---|---|---|---|---|
| Mean | 56 | 183 | 181 | 153 | 126 | 95 | 84 | 55 | 60 | 58 | 64 | 51 |
| Std | 14 | 44 | 7 | 14 | 15 | 17 | 18 | 8 | 2 | 6 | 3 | 5 |
| N | 10 | 10 | 10 | 10 | 15 | 29 | 15 | 7 | 6 | 10 | 3 | 9 |
| DISTRIBUTION NORMALITY TESTS | | | | | | | | | | | | |
| Shapiro-Wilk | ns | *** | ns | ns | * | *** | ns | ns | ns | ns | *** | ns |
| Kolmogorov-Smirnov | ns | * | ns | ns | * | *** | ns | ns | ns | * | | ns |
| Cramer-von Mises | ns | ** | ns | ns | * | *** | ns | | | ns | | ns |
| Anderson-Darling | ns | *** | ns | ns | * | *** | ns | | | ns | | ns |
| ANOVA | | | | | | | | | | | | |
| Year Effect | | | | | *** | *** | ns | ns | | ns | | |
| Yearly means | | | | | | | | | | | | |
| 2006 | | | | | | | | | | 54 a | 64 | 51 |
| 2007 | 56 | 183 | 181 | 153 | 138 a | 106 a | 75 a | 51 a | | | | |
| 2008 | | | | | 112 b | 72 b | 92 a | 62 a | 60 | 60 a | | |
