## Supplementary Figures and Tables for "Consequence and further analysis of the minor role of daylength in the floral initiation of sorghum crops"

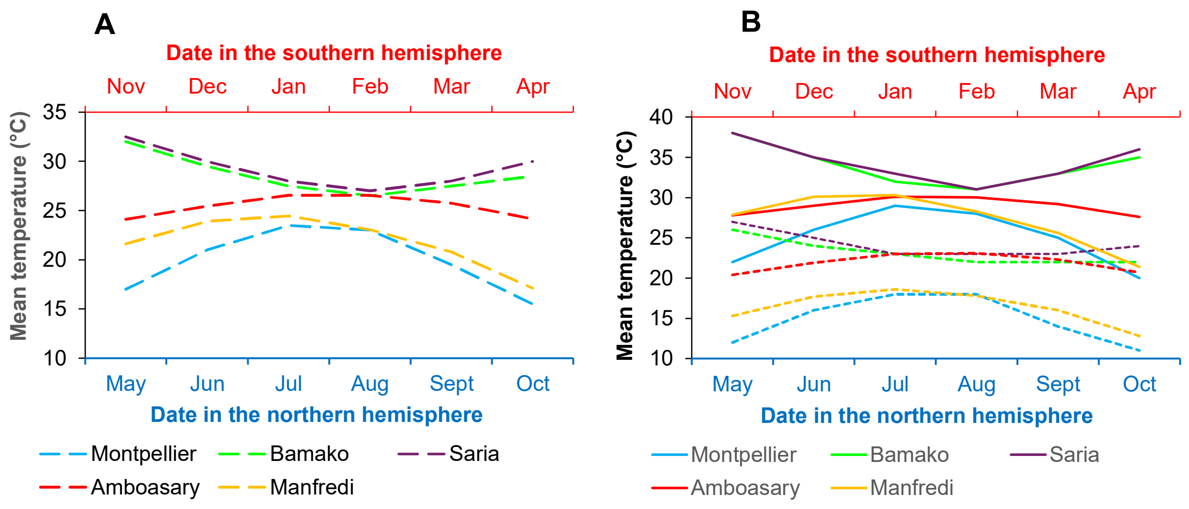


**Fig. S1:** Monthly mean of **A**) the daily average temperature (dashed ligne), and **B**) the daily maximum (solid line) and minimum (dotted line) temperatures during the cropping season in five locations from both the northern and southern hemispheres where the duration to flowering of sorghum crops is compared.

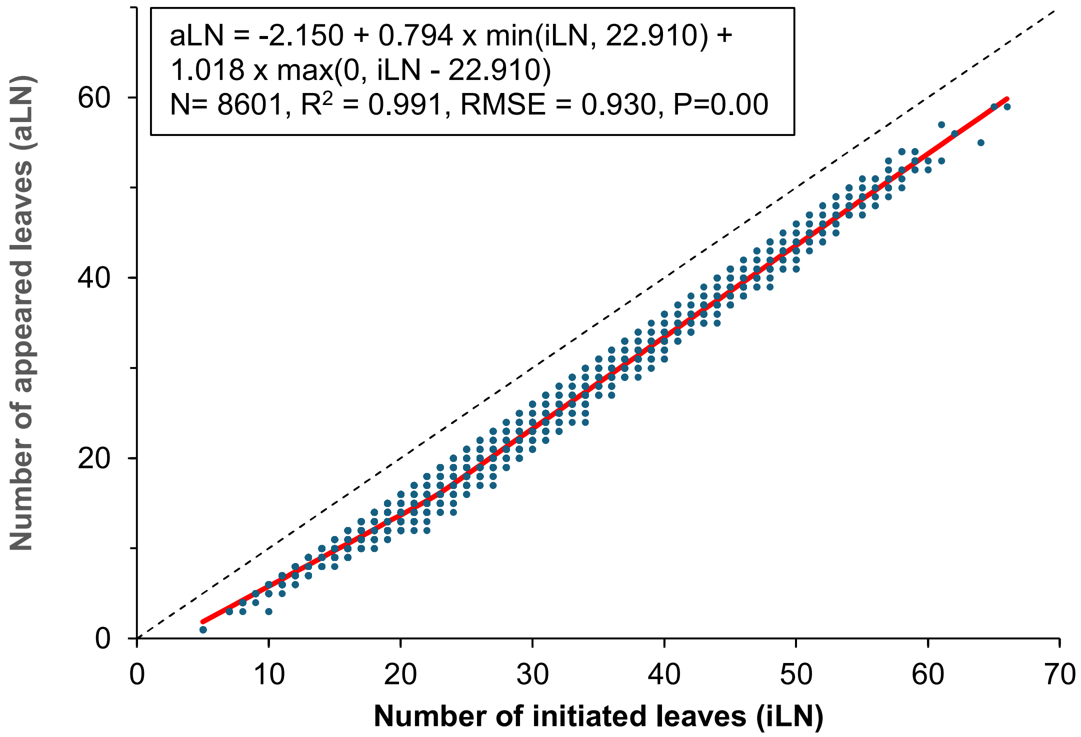


**Fig. S2:** Segmented linear regression of appeared leaf number (aLN) on initiated leaf number (iLN), using all observations from the development data of 28 sorghum varieties, where iLN > aLN by at least 3. Other observations were generally recorded after panicle initiation, by which point no further leaves were initiated. The number of leaves developing inside the leaf sheaths increased from four at plant emergence to an average of seven by the time the 23rd leaf was initiated, which coincided with the appearance of the 16th leaf. Thereafter, the difference of seven leaves decreased slowly as more leaves were produced.

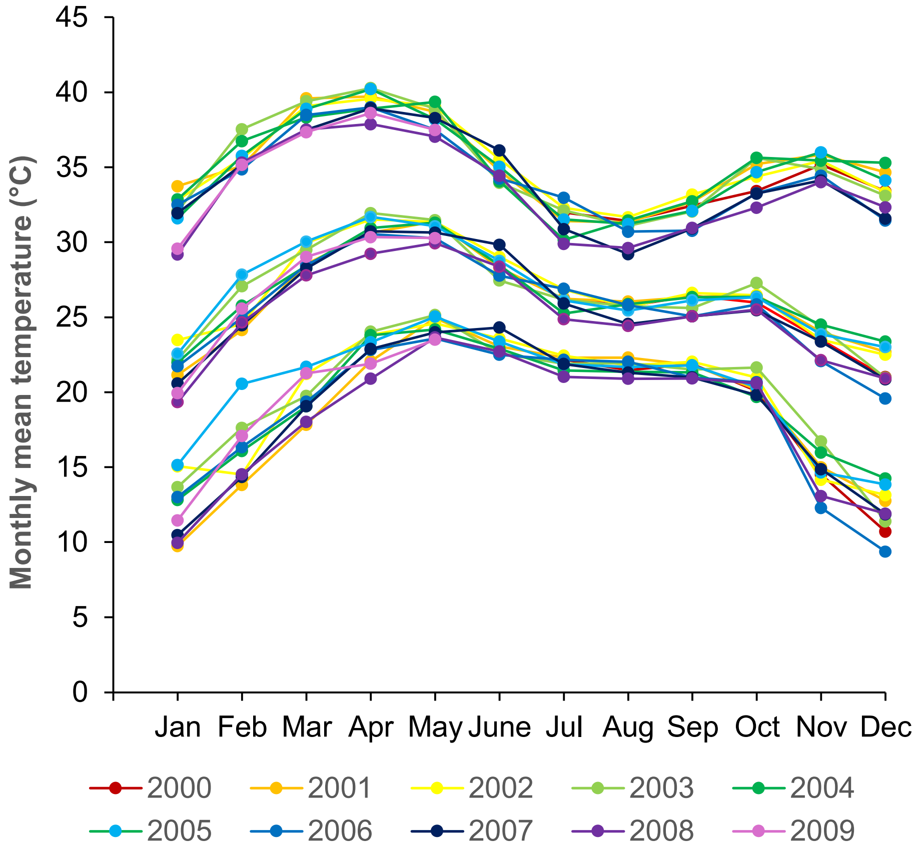


**Fig. S3:** Monthly maximum, average, and minimum temperatures recorded in Samanko, Mali, from July 2000 to May 2009.

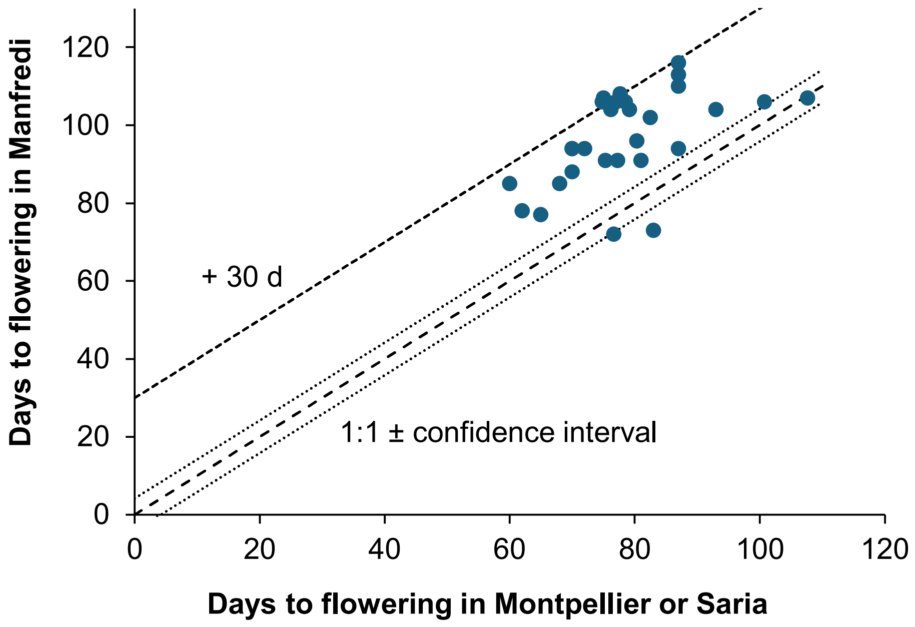


**Fig. S4:** Comparison between the duration to flowering of 27 cultivars observed in the northern hemisphere in either Montpellier (France), Saria (Burkina Faso), and in the southern hemisphere in Manfredi (Argentina).

**Table S1:** List, origin, and durations to heading of 74 early to mid-early sorghum inbred lines and progenies developed in the Northern Hemisphere and 3 cultivars from Southern Africa and observed in a variety adaptation trial in southern Madagascar in 2020-2021. The trial was sown on 14 Nov and some lines were resown on 5 Dec and the duration to heading was reported accordingly. The reference heading date was observed in the Northern Hemisphere.

| | | | | Duration to heading (d) | | | |
|---|---|---|---|---|---|---|---|
| | | | | Sowing date | | Reference sowing | |
| Variety/Line name | Origin | Seed Bank | Plant height | 14 Nov | 5 Dec | May | July |
| 06-SB-F5DT-15 | Mali | IER | 182 | >150 | | | 65 |
| Darrel Ken | Mali | IER | 245 | >150 | | | 45 |
| Dougouyiriwa | Mali | IER | 240 | 103 | | | 70 |
| Saba-Nafanté | Mali | IER | 184 | 109 | | | 62 |
| Sangatigui | Mali | IER | 253 | >150 | | | 55 |
| Seguifa | Mali | IER | 100 | 134 | | | 55 |
| F4 SSM10-13/7-1 | Mali | ICRISAT | | >150 | | | 84 |
| F4 SSM10-14/1-1 | Mali | ICRISAT | | >150 | | | 91 |
| F4 SSM10-21/10-1 | Mali | ICRISAT | | >150 | | | 76 |
| F4 SSM10-21/4-1 tan | Mali | ICRISAT | | >150 | | | 82 |
| F4 SSM10-21/8-1 | Mali | ICRISAT | | >150 | | | 69 |
| Soubatimi | Mali | ICRISAT | | >150 | | | 65 |
| Soumba | Mali | ICRISAT | | >150 | | | 83 |
| GPNDC2 S1.0-AbdKon-16-1-1-2-1-SB-2-SB | | ICRISAT | 193 | | 138 | | 84 |
| ICSX 1560002-6-10-1-SB-1 | | ICRISAT | 205 | | 138 | | 85 |
| ICSX 1560003-12-4-SB3-1 | | ICRISAT | | >150 | | | 85 |
| ICSX 1560003-12-5-1-SB-1-SB | | ICRISAT | 148 | | 138 | | 82 |
| LocBl-2-19-2-RB5-SB-1-SB | | ICRISAT | 363 | 138 | 138 | | 85 |
| LocBl-6-1-1-RB2-SB-1-SB | | ICRISAT | 193 | 134 | | | 84 |
| PGD_S3_SamKon-29-VS4.2-2-2-SB-1-SB | | ICRISAT | | | 146 | | 84 |
| PGND12C2-S1-284-1-7-2-SB-1-SB | | ICRISAT | 247 | | 138 | | 84 |
| SC 1033 (IS 11894c) | Ethiopia | CIRAD | | 68 | | 93 | 60 |
| SC 135 (IS 12626c) | Ethiopia | CIRAD | | 61 | 55 | 84 | 83 |
| SC 982 (IS 1256c) | Ethiopia | CIRAD | | 88 | 40 | 85 | 65 |
| BF201 | France | CIRAD | 100 | 57 | 53 | 58 | |
| IS 18164 | Lebanon | CIRAD | | 77 | | 62 | 67 |
| IRAT 151 | Mali | CIRAD | 100 | 58 | 53 | 68 | |
| CIR-13/15Z-1-1-1 | Nicaragua | CIRAD | 120 | 67 | | 74 | |
| CIR-13/8Z-1-1-V1 | Nicaragua | CIRAD | 80 | 58 | 54 | 76 | |
| CIR-9/32Z-4-1-1 | Nicaragua | CIRAD | 80 | 72 | | 74 | |
| 3241 | Niger | CIRAD | 98 | >150 | | 76 | 71 |
| SC 525 (IS4832c) | Nigeria | CIRAD | | 51 | 55 | 67 | 54 |
| Persis-5 | Russia | CIRAD | 130 | 75 | | 78 | |
| Russia Sugar-2 | Russia | CIRAD | 180 | 61 | | 82 | |
| SKI-6 | Russia | CIRAD | 200 | 55 | | 82 | |
| IS 2262 | Soudan | CIRAD | | 109 | | 87 | 77 |
| PI 308433 | South Africa | CIRAD | 100 | 77 | | 77 | 72 |
| SC 805 (IS 2732c) | Uganda | CIRAD | | 88 | 58 | 86 | 83 |
| SC 673 (IS 2540c) | Zimbabwe | CIRAD | | 94 | 53 | 75 | 55 |

| | | | Duration to heading (d) | |
|---|---|---|---|---|
| | | | Sowing | Reference |
| Breeding line | Seeds | Plant height | 14 Nov | May |
| SSM10-21/8 | CIRAD | 160 | 138 | 80 |
| SSM10-14 | CIRAD | 175 | 135 | 80 |
| SSM10-21/10 | CIRAD | 180 | 122 | 80 |
| SSM10-13 | CIRAD | 170 | 135 | 80 |
| SSM10-21/4 | CIRAD | 190 | 135 | 80 |
| IRAT204-1 | CIRAD | 100 | 138 | 80 |
| IRAT204-2 | CIRAD | 110 | 135 | 80 |
| IRAT204-3 | CIRAD | 100 | 122 | 80 |
| IRAT204-4 | CIRAD | 100 | 135 | 80 |
| IRAT204-5 | CIRAD | 110 | 135 | 80 |
| IRAT204-6 | CIRAD | 120 | 49 | 60 |
| IRAT204-7 | CIRAD | 160 | 57 | 60 |
| IRAT204-8 | CIRAD | 170 | 57 | 60 |
| IRAT204-9 | CIRAD | 100 | | 60 |
| IRAT204-10 | CIRAD | 100 | 59 | 60 |
| IRAT204-11 | CIRAD | 100 | 61 | 60 |
| IRAT204-12 | CIRAD | 90 | 55 | 60 |
| IRAT204-13 | CIRAD | 100 | 59 | 60 |
| IRAT204-14 | CIRAD | 150 | 67 | 60 |
| IRAT204-15 | CIRAD | 120 | | 60 |
| IRAT204-16 | CIRAD | 100 | | 60 |
| IRAT204-17 | CIRAD | 110 | 61 | 60 |
| IRAT204-18 | CIRAD | 190 | 61 | 60 |
| IRAT204-19 | CIRAD | 120 | 55 | 60 |
| IRAT204-20 | CIRAD | 100 | 57 | 60 |
| IRAT204-21 | CIRAD | 100 | 59 | 60 |
| AD-6-1 | CIRAD | 110 | 61 | 60 |
| AD-6-2 | CIRAD | 214 | 55 | 60 |
| AD-6-3 | CIRAD | 170 | 59 | 60 |
| AD-6-4 | CIRAD | 160 | 70 | 60 |
| AD-6-5 | CIRAD | 150 | 57 | 60 |
| AD-6-6 | CIRAD | 150 | 55 | 70 |
| AD-6-7 | CIRAD | 170 | 61 | 70 |
| AD-6-8 | CIRAD | 260 | 62 | 70 |
| AD-6-9 | CIRAD | 230 | 59 | 70 |
| AD-6-10 | CIRAD | 230 | 81 | 70 |
| AD-6-11 | CIRAD | 190 | 61 | 70 |
| PCR-2 | CIRAD | 190 | 54 | 70 |

**Table S2:** List of the 28 sorghum varieties monthly sown in Samanko, Mali.

| Genotype | Geographic origin | Race | Number of months |
|---|---|---|---|
| Blanc Augaradeba | Nigeria | Guinea | 29 |
| CGM 19 | Mali | Guinea-caudatum | 22 |
| CS 02 Mdg | Madagascar | Guinea margaritiferum | 29 |
| CSM 335 | Mali | Guinea | 76 |
| CSM 63E | Mali | Guinea | 24 |
| Farafara 17 | Nigeria | Guinea | 29 |
| IBS 19 | Tanzanie | Guinea | 24 |
| IBS 30 | Tanzanie | Guinea | 24 |
| IBS 40 | Tanzanie | Guinea | 24 |
| IBS 582 | Mozambique | Guinea | 24 |
| IRAT 174 | Burkina Faso | Kafir-durra | 50 |
| IRAT 204 | Senegal | Caudatum | 28 |
| IS 11026 | Ethiopia | Durra | 14 |
| IS 12169 | Ethiopia | Bicolor | 14 |
| IS 15401 | Cameroon | Guinea-caudatum | 24 |
| IS 6351 | India | Durra | 14 |
| Kaura D-12 | Nigeria | Durra-caudatum | 29 |
| Kendé Bla | Mali | Guinea margaritiferum | 23 |
| Macia | Zimbabwe | Caudatum | 24 |
| Mtama | Kenya | Caudatum | 24 |
| Ouéni | Mali | Guinea | 24 |
| Sariaso 10 | Burkina Faso | Caudatum | 50 |
| Short Kaura | Nigeria | Durra-caudatum | 24 |
| Sima | Zimbabwe | Guinea-caudatum | 24 |
| Souroukoukou | Mali | Caudatum | 22 |
| SSM 12 | Cameroon | Durra | 25 |
| SSM 1592 | Chad | Durra | 24 |
| SSM 29 | Cameroon | Durra | 17 |

**Table S3:** Comparison between the number of days to flowering of 27 cultivars sown either in the southern hemisphere (Manfredi, Argentina) or in the northern hemisphere (Montpellier, France; Saria, Burkina Faso; and Cali, Colombia).

| | | Manfredi | Montpellier | | | | | | | Saria | Cali |
|---|---|---|---|---|---|---|---|---|---|---|---|
| | Sowing date | 07/12 2011 | 16/05 2008 | 14/05 2009 | 20/05 2010 | 17/05 2011 | 14/05 2012 | 04/05 2014 | 13/05 2015 | Mid-June | 29/11 2007 |
| Cultivars | Origin | | | | | | | | | | |
| CIR-1/OG2-4G-1G-M-M | Nicaragua | 110 | | | 97 | | | | | | 66 |
| Cirad 440 | Burkina Faso | 94 | | | | | 89 | | | 75 | |
| Cirad 492 | Burkina Faso | 94 | | | | | | | | 80 | 52 |
| BF 94-6/11-1K-1K | Burkina Faso | 78 | | | | | | | | 72 | 64 |
| Sorgo blanco alto | Nicaragua | 104 | | | | | 103 | | | | 61 |
| IS 6193 | India | 108 | 88 | | | 90 | 85 | | | | |
| IS 29375 | Lesotho | 106 | 86 | 91 | | | | | | | |
| IS 18461 | India | 91 | | | 91 | | | | | | 68 |
| IS 2787 | Kenya | 91 | 94 | 85 | | | 80 | 90 | | | |
| IS 5972 | India | 113 | 104 | 90 | | | | | | | |
| IS 15752 | Cameroon | 91 | 87 | 82 | | | 87 | | | | |
| IS 16044 | Cameroon | 96 | 94 | 90 | | | 87 | | | | |
| IS 22332 | Bostwana | 116 | 99 | 94 | 98 | | 94 | 100 | | | |
| IS 26833 | Sudan | 106 | 113 | 108 | 111 | | 111 | | | | |
| IS 28409 | Yemen | 104 | 88 | 93 | 82 | 94 | 88 | 90 | | | |
| IS 32569 | Somalia | 107 | 127 | 110 | | | 116 | | | | |
| IS 19358 | Sudan | 104 | | 81 | 78 | 91 | 88 | 93 | | | |
| IS 18164 | Lebanon | 102 | | 87 | | 98 | 89 | 96 | | | 57 |
| SSM 264 | Burkina Faso | 94 | | | | | | | 97 | | 46 |
| SSM 1264 | Eswatini | 94 | | | | | | | | 70 | 70 |
| SSM 1279 | Nigeria | 85 | | | | | | | | 60 | 60 |
| SSM 1571 | Cameroon | 85 | | | | | | | | 68 | 68 |
| BMA 18-2-1(2) | France | 73 | | | | | 93 | | | | |
| BMA 12-2-1 | France | 77 | | | 75 | | | | | | |
| IS 3780 | China | 106 | 87 | 80 | 75 | 88 | 90 | 89 | | | |
| IS 30435 | China | 88 | | 75 | 73 | 85 | 84 | 83 | | | |
| Russia Sugar-2 | Russia | 72 | | | 89 | 87 | 84 | | | | |

**Table S5:** Optimised parameters of the four photoperiodic responses for each variety grown in Samanko, the number of plants (n) and the resulting RMSE computed by plant for the model using only dSR and dSS (noDL). PIP is used for Kt1 and QL1, and PIP2 for Kt2, QL2 and QL2s. The RMSE resulting from the initial model in which daylength (+DL) was still used in the Kt1 and Kt2 equations are given together with the recomputed RMSE for Photoperiodism Model 2.0 (PP2.0). The RMSE computed for all 28 varieties together is shown on the last line.

| Variety | Kt1 | | | | | | | Kt2 | | | | QL1 | | | QL2 | | | | | | n | RMSE | | |
|---|---|---|---|---|---|---|---|---|---|---|---|---|---|---|---|---|---|---|---|---|---|---|---|---|
| | PIP1 | dsr11 | dss11 | B11 | dsr12 | dss12 | B12 | PIP2 | dsr2 | dss2 | B2 | Bcst1 | Bdsr1 | Bdss1 | Bcst2 | Bdsr2 | Bdss2 | Bcst2s | Bdsr2s | Bdss2s | | noDL | +DL | PP2.0 |
| Blanc Augaradeba | 19.16 | 1.81 | 1.01 | 88.40 | -0.20 | -0.09 | 20.00 | 6.42 | -0.06 | 0.07 | 41.29 | -39.23 | -0.27 | -0.75 | | | | | | | 284 | 14.25 | 12.90 | 13.63 |
| CGM 19 | 17.09 | 0.40 | 0.86 | 54.45 | 0.05 | 0.45 | 25.17 | 12.11 | -0.13 | 0.57 | 43.98 | -2.17 | -0.70 | 0.41 | 31.70 | -0.10 | -0.61 | | | | 190 | 6.12 | 6.77 | 21.69 |
| CS02 Mdg | 22.80 | 1.55 | 1.01 | 75.30 | -0.21 | 0.05 | 18.73 | 22.98 | -0.66 | -0.26 | 68.65 | -22.49 | -0.70 | -0.58 | | | | | | | 289 | 11.06 | 11.79 | 14.07 |
| CSM 335 | 19.30 | 0.17 | 1.03 | 59.98 | -0.26 | 0.09 | 11.94 | 18.70 | -0.07 | 0.33 | 25.07 | -2.79 | -0.77 | -0.16 | -27.30 | 0.43 | -0.30 | | | | 668 | 9.40 | 9.60 | 20.55 |
| CSM 63-E | 19.06 | -0.20 | 0.12 | 14.55 | -0.41 | 0.12 | 10.87 | 17.65 | -0.20 | 0.64 | 34.89 | | | | -28.38 | -1.16 | 2.99 | | | | 237 | 5.36 | 5.41 | 13.34 |
| Farafara 17 | 20.43 | 1.53 | 1.09 | 78.38 | -0.20 | 0.07 | 14.42 | 11.26 | -0.10 | 0.43 | 39.85 | -25.53 | -0.68 | -0.18 | | | | | | | 276 | 11.99 | 11.88 | 18.01 |
| IBS 19 | 20.09 | 1.44 | 1.05 | 83.49 | -0.39 | -0.09 | 24.90 | 7.41 | -1.32 | 0.82 | 88.26 | -31.23 | -0.49 | -0.37 | 6.04 | 0.37 | 0.22 | 6.39 | 0.73 | -0.38 | 199 | 16.97 | 16.92 | 27.27 |
| IBS 30 | 41.30 | 2.24 | 0.92 | 71.24 | -0.62 | 0.37 | 19.58 | 41.62 | -1.47 | 0.97 | 39.27 | -34.02 | -0.41 | 0.15 | -39.48 | -0.36 | 1.36 | -31.62 | -2.62 | 5.29 | 200 | 25.14 | 12.61 | 28.90 |
| IBS 40 | 17.59 | 0.91 | 0.82 | 65.19 | -0.05 | 0.12 | 24.53 | 28.28 | -0.56 | -1.16 | 112.81 | -33.76 | -0.32 | -0.07 | -0.41 | 0.30 | 0.40 | -0.15 | 0.50 | -0.12 | 200 | 13.55 | 13.53 | 27.54 |
| IBS 582 | 21.74 | 1.60 | 1.09 | 78.56 | -0.40 | 0.01 | 22.03 | 5.78 | 0.83 | 1.63 | 91.55 | -32.28 | -0.36 | -0.74 | -22.71 | -0.06 | 1.08 | -17.25 | 0.72 | 0.77 | 196 | 14.25 | 13.09 | 32.44 |
| IRAT 174 | 22.20 | 0.89 | 0.43 | 37.84 | -0.20 | 0.07 | 4.94 | 11.98 | 0.18 | 0.56 | 19.69 | -2.67 | -1.18 | -0.11 | -6.73 | 0.63 | -0.91 | | | | 461 | 5.59 | 6.27 | 9.50 |
| IRAT 204 | 12.45 | 0.31 | -0.16 | 20.14 | 0.02 | -0.05 | 11.19 | 7.00 | | | | | | | | | | | | | 268 | 5.57 | 6.18 | 6.65 |
| IS 11026 | 29.23 | 0.23 | 0.31 | 38.10 | -0.69 | 0.02 | 24.72 | 23.12 | -0.18 | -0.27 | 61.09 | 4.24 | -0.82 | -0.29 | -26.41 | 3.42 | -6.37 | | | | 115 | 10.02 | 9.46 | 16.14 |
| IS 12169 | 23.07 | 1.16 | 0.93 | 65.91 | -0.40 | 0.13 | 15.53 | 5.00 | -0.84 | 0.72 | 49.21 | -22.56 | -0.71 | 0.95 | -6.35 | -1.03 | 1.00 | -6.37 | -0.56 | 0.95 | 118 | 11.07 | 11.22 | 30.77 |
| IS 15401 | 4.21 | 0.17 | -0.35 | 73.64 | -0.16 | 0.06 | 31.59 | 21.78 | | | | -39.62 | -0.18 | -0.09 | | | | | | | 240 | 11.47 | 11.26 | 15.89 |
| IS 6351 | 19.40 | -0.53 | 0.65 | 59.89 | -0.39 | 0.32 | 29.78 | 7.00 | | | | | | | -10.67 | -0.40 | -0.05 | | | | 122 | 6.94 | 6.72 | 8.45 |
| Kaura D-12 | 13.05 | 1.15 | 1.05 | 78.09 | -0.40 | 0.18 | 17.09 | 19.48 | -0.62 | 0.29 | 30.58 | -120.97 | -2.72 | 4.82 | | | | | | | 281 | 12.84 | 12.72 | 18.17 |
| Kende Bla | 18.29 | -0.10 | 0.44 | 27.74 | -0.12 | 0.07 | 13.45 | 19.00 | | | | -31.62 | -1.32 | -1.99 | | | | | | | 203 | 5.50 | 5.87 | 12.01 |
| Macia | 18.91 | 0.08 | -0.15 | 17.14 | -0.10 | -0.18 | 11.53 | 15.74 | -0.04 | 0.26 | 18.53 | | | | | | | | | | 214 | 5.14 | 5.29 | 6.90 |
| Mtama | 20.32 | -0.01 | 0.15 | 28.09 | -0.21 | 0.21 | 14.16 | 16.02 | 0.69 | -0.74 | 50.85 | | | | 36.11 | 0.79 | -1.30 | | | | 219 | 5.12 | 5.57 | 6.74 |
| Oueni | 19.32 | -0.05 | 0.59 | 39.03 | -0.38 | 0.18 | 23.38 | 22.00 | | | | 3.64 | -0.51 | 0.63 | 19.06 | 0.63 | -0.55 | | | | 222 | 10.67 | 10.96 | 15.97 |
| Sariaso 10 | 17.25 | 0.10 | 0.35 | 35.84 | -0.41 | 0.14 | 18.09 | 13.41 | -0.27 | 0.29 | 25.98 | | | | 5.35 | 0.37 | -0.56 | | | | 454 | 6.94 | 7.21 | 9.54 |
| Short Kaura | 19.33 | 1.61 | 1.03 | 99.15 | -0.88 | -0.11 | 33.93 | 14.69 | -0.86 | -0.17 | 59.77 | -31.92 | -0.57 | -1.23 | | | | | | | 237 | 18.28 | 18.09 | 16.88 |
| Sima | 17.46 | -0.03 | 0.22 | 33.63 | -0.29 | 0.08 | 13.07 | 15.00 | -0.19 | 0.39 | 25.52 | | | | | | | | | | 213 | 5.36 | 5.44 | 6.85 |
| Souroukoukou | 10.91 | 1.92 | -0.01 | 107.88 | -0.44 | -0.06 | 28.09 | 15.26 | -0.27 | 0.27 | 34.68 | -39.24 | -0.21 | -0.90 | | | | | | | 213 | 12.98 | 12.79 | 15.81 |
| SSM12 | 18.43 | -2.06 | 0.55 | 72.22 | 0.26 | 0.06 | 23.66 | 22.66 | -0.53 | 0.83 | 41.86 | -9.70 | -0.50 | -3.37 | | | | | | | 184 | 14.13 | 16.38 | 17.11 |
| SSM1592 | 10.80 | 1.21 | 0.57 | 78.88 | -0.41 | 0.21 | 40.50 | 20.37 | -0.83 | 0.89 | 57.62 | -41.28 | -0.31 | 0.00 | | | | | | | 179 | 13.87 | 13.47 | 22.39 |
| SSM29 | 20.49 | 1.87 | 1.12 | 95.55 | 0.06 | -0.10 | 37.56 | 21.00 | | | | -40.38 | -0.05 | -0.40 | | | | | | | 124 | 13.81 | 13.08 | 17.99 |
| 28 varieties | | | | | | | | | | | | | | | | | | | | | 6806 | 11.37 | 10.77 | 17.59 |

**Table S6:** Optimised parameters of the four photoperiodic responses for each variety grown in Samanko, the number of plants (n) and the resulting RMSE computed by plant for the model using DL, dSR and dSS in the Kt1 and Kt2 equations and only dSR and dSS in the QL equations (DL). PIP is used for Kt1 and QL1, and PIP2 for Kt2, QL2 and QL2s. The RMSE computed for all 28 varieties together is shown on the last line.

| Variety | Kt1 | | | | | | Kt2 | | | | | QL1 | | | QL2 | | | | | | n | RMSE DL |
|---|---|---|---|---|---|---|---|---|---|---|---|---|---|---|---|---|---|---|---|---|---|---|
| | PIP1 | DLmin1 | DLmax | slp1 | dsr1 | dss1 | PIP2 | DLmin2 | Slp2 | dsr2 | dss2 | Bcst1 | BdSR1 | BdSS1 | Bcst2 | Bdsr2 | Bdss2 | Bcst2s | Bdsr2s | Bdss2s | | |
| Blanc Augaradeba | 15.71 | 11.33 | 16.07 | 64.04 | 0.13 | 0.28 | 30.60 | 11.50 | 83.14 | 0.34 | 0.35 | -129.26 | -2.15 | 4.07 | | | | | | | 284 | 12.90 |
| CGM 19 | 21.30 | 10.48 | 14.86 | 16.94 | 0.02 | 0.40 | 23.91 | 10.62 | 29.24 | 0.04 | 0.71 | -2.30 | -0.71 | 0.53 | 31.70 | -0.10 | -0.61 | | | | 190 | 6.77 |
| CS02 Mdg | 14.75 | 11.25 | 19.17 | 57.46 | 0.21 | 0.45 | 14.74 | 10.65 | 62.57 | -0.17 | 0.33 | -18.40 | -0.53 | -0.89 | | | | | | | 289 | 11.79 |
| CSM 335 | 15.28 | 11.16 | 15.90 | 34.19 | -0.09 | 0.34 | 25.31 | 11.35 | 57.71 | 0.22 | 0.59 | -2.79 | -0.77 | -0.16 | -31.24 | 0.38 | -0.16 | | | | 668 | 9.60 |
| CSM 63-E | 20.00 | 9.33 | 15.37 | 4.00 | -0.35 | 0.08 | 25.10 | 9.52 | 12.03 | -0.18 | 0.69 | | | | -28.84 | -1.10 | 2.74 | | | | 237 | 5.41 |
| Farafara 17 | 17.57 | 11.41 | 16.27 | 57.18 | 0.13 | 0.35 | 18.44 | 11.18 | 65.46 | 0.27 | 0.74 | -25.67 | -0.67 | -0.29 | | | | | | | 276 | 11.88 |
| IBS 19 | 12.09 | 11.04 | 15.90 | 45.89 | 0.02 | 0.13 | 2.81 | 10.57 | 73.73 | -0.35 | 1.11 | -31.87 | -0.44 | -0.33 | 6.04 | 0.37 | 0.22 | 6.44 | 0.67 | -0.30 | 199 | 16.92 |
| IBS 30 | 25.05 | 10.70 | 16.66 | 36.56 | -0.33 | 0.65 | 26.33 | 10.64 | 42.55 | -0.66 | 1.99 | -35.47 | -0.42 | -0.07 | -39.48 | -0.36 | 1.36 | -35.36 | -0.25 | 1.26 | 200 | 12.61 |
| IBS 40 | 20.35 | 11.11 | 15.38 | 39.49 | 0.45 | 0.36 | 30.94 | 7.70 | 27.09 | -0.13 | -1.06 | -34.78 | -0.33 | 0.05 | -0.41 | 0.30 | 0.40 | -0.06 | 0.49 | -0.32 | 200 | 13.53 |
| IBS 582 | 17.04 | 11.10 | 16.49 | 41.20 | 0.21 | 0.20 | 7.88 | 10.02 | 45.88 | 0.87 | 1.55 | -28.48 | -0.27 | -0.76 | -22.71 | -0.06 | 1.08 | -21.91 | 0.27 | 1.08 | 196 | 13.09 |
| IRAT 174 | 0.69 | 10.94 | 30.21 | 37.66 | 0.56 | 0.59 | 15.78 | 13.62 | 54.78 | 0.20 | 1.07 | -3.08 | -0.75 | -0.89 | -7.36 | 0.46 | -1.12 | | | | 461 | 6.27 |
| IRAT 204 | 7.27 | 10.25 | 14.14 | 10.01 | 0.13 | -0.08 | 7.00 | 10.30 | 17.19 | 0.18 | 0.05 | | | | | | | | | | 268 | 6.18 |
| IS 11026 | 22.85 | 11.21 | 12.18 | 46.99 | 0.13 | 0.30 | 23.12 | 10.55 | 29.18 | 0.07 | 0.78 | 5.25 | -1.20 | -0.52 | -26.41 | 3.42 | -6.37 | | | | 115 | 9.46 |
| IS 12169 | 16.30 | 11.31 | 14.71 | 47.84 | 0.08 | 0.30 | 17.97 | 11.03 | 52.02 | -0.48 | 0.85 | -25.66 | -0.78 | 0.69 | -6.35 | -1.03 | 1.00 | -6.16 | -0.67 | 0.70 | 118 | 11.22 |
| IS 15401 | 20.13 | 11.42 | 19.06 | 49.32 | 0.19 | 0.16 | 21.78 | 11.47 | | | | -40.02 | -0.18 | -0.08 | | | | | | | 240 | 11.26 |
| IS 6351 | 8.05 | 10.59 | 14.62 | 32.58 | -0.57 | 0.40 | 7.00 | 10.80 | | | | | | | -10.67 | -0.40 | -0.05 | | | | 122 | 6.72 |
| Kaura D-12 | 12.55 | 11.38 | 16.87 | 56.92 | 0.00 | 0.46 | 19.30 | 11.47 | 84.87 | 0.16 | 0.70 | -117.70 | -2.68 | 4.63 | | | | | | | 281 | 12.72 |
| Kende Bla | 18.87 | 11.00 | 14.76 | 16.42 | -0.09 | 0.27 | 19.00 | 10.99 | | | | -33.02 | -1.27 | -1.70 | | | | | | | 203 | 5.87 |
| Macia | 16.19 | 9.99 | 14.43 | 7.55 | -0.05 | -0.21 | 10.99 | 10.25 | 15.61 | 0.07 | 0.33 | | | | | | | | | | 214 | 5.29 |
| Mtama | 16.69 | 10.18 | 14.01 | 12.38 | -0.15 | 0.14 | 16.87 | 10.33 | 14.68 | 0.66 | 0.41 | | | | 34.60 | 0.79 | -0.97 | | | | 219 | 5.57 |
| Oueni | 22.12 | 10.82 | 14.07 | 20.51 | -0.21 | 0.47 | 22.00 | 10.81 | | | | 2.79 | -0.71 | 0.66 | 18.39 | 0.79 | -0.60 | | | | 222 | 10.96 |
| Sariaso 10 | 19.07 | 10.87 | 13.57 | 19.67 | -0.15 | 0.22 | 18.06 | 10.90 | 29.05 | -0.15 | 0.43 | | | | 4.51 | 0.53 | -0.57 | | | | 454 | 7.21 |
| Short Kaura | 16.57 | 11.07 | 17.66 | 55.81 | -0.35 | 0.18 | 14.36 | 10.73 | 63.95 | -0.37 | 0.15 | -34.39 | -0.50 | -0.45 | | | | | | | 237 | 18.09 |
| Sima | 15.99 | 11.04 | 13.36 | 21.90 | -0.22 | 0.12 | 15.52 | 10.82 | 33.38 | -0.06 | 0.44 | | | | | | | | | | 213 | 5.44 |
| Souroukoukou | 14.12 | 11.38 | 17.71 | 83.65 | 0.16 | 0.32 | 14.75 | 11.34 | 104.34 | 0.19 | 0.87 | -39.24 | -0.22 | -0.95 | | | | | | | 213 | 12.79 |
| SSM12 | 6.64 | 10.73 | 16.16 | 35.53 | 0.48 | 0.21 | 19.71 | 10.84 | 71.51 | -0.54 | 0.97 | -11.04 | -0.24 | -1.94 | | | | | | | 184 | 16.38 |
| SSM1592 | 28.08 | 11.04 | 15.86 | 37.56 | -0.05 | 0.22 | 29.02 | 11.10 | 61.23 | -0.15 | 1.28 | -43.57 | -0.46 | -0.16 | | | | | | | 179 | 13.47 |
| SSM29 | 20.70 | 10.74 | 14.08 | 38.07 | 0.32 | 0.08 | 21.00 | 10.72 | | | | -44.54 | -0.14 | -0.23 | | | | | | | 124 | 13.08 |
| 28 varieties | | | | | | | | | | | | | | | | | | | | | 6807 | 10.77 |

**Table S7:** Optimised parameters of the four photoperiodic responses for three varieties grown in Samanko and in Montpellier, the number of year*month data (n) and the resulting RMSE computed on the year*month means for the model using only dSR and dSS (noDL). The RMSE for Photoperiodism Model 2.0 (PP2.0), computed on all year*month data is given.

| | Kt1 | | | | | | | Kt2 | | | QL1 | | | | QL2 | | | RMSE | | | |
|---|---|---|---|---|---|---|---|---|---|---|---|---|---|---|---|---|---|---|---|---|---|
| Variety | PIP | dsr11 | dss11 | B11 | dsr12 | dss12 | B12 | dsr2 | dss2 | B2 | Bcst1 | Bdsr1 | Bdss1 | dl1 | Bcst2 | BdSR2 | BdSS2 | n | noDL | n | PP2.0 |
| CSM 335 | 21.26 | 1.00 | 0.84 | 51.73 | -0.19 | 0.07 | 11.66 | -0.15 | 0.33 | 24.62 | -2.65 | -0.72 | -0.18 | -1.52 | -28.15 | 0.40 | -0.21 | 81 | 8.17 | 70 | 14.29 |
| Sariaso 10 | 18.32 | 0.08 | 0.33 | 35.24 | -0.35 | 0.20 | 17.29 | -0.36 | 0.25 | 23.82 | | | | | 4.90 | 0.57 | -0.63 | 61 | 4.29 | 57 | 9.71 |
| Souroukoukou | 18.91 | 2.15 | 1.53 | 99.16 | -0.18 | -0.05 | 24.28 | -0.18 | 0.20 | 31.64 | -41.71 | -0.30 | -1.49 | 0 | | | | 25 | 7.36 | 25 | 10.85 |
